# Understanding the Separation of Timescales in *Rhodococcus erythropolis* Proteasome Core Particle Assembly

**DOI:** 10.1101/2022.02.04.479176

**Authors:** Pushpa Itagi, Anupama Kante, Leonila Lagunes, Eric J. Deeds

## Abstract

The 20S proteasome Core Particle (CP) is a molecular machine that is a key component of cellular protein degradation pathways. Like other molecular machines, it is not synthesized in an active form, but rather as a set of subunits that assemble into a functional complex. The CP is conserved across all domains of life and is composed of 28 subunits, 14 α and 14 β, arranged in four stacked 7-member rings (α_7_β_7_β_7_α_7_). While details of CP assembly vary across species, the final step in the assembly process is universally conserved: two half proteasomes (HP: α_7_β_7_) dimerize to form the CP. In the bacterium *Rhodococcus erythropolis*, experiments have shown that the formation of the HP is completed within minutes, while the dimerization process takes hours. The N-terminal propeptide of the β subunit, which is autocatalytically cleaved off after CP formation, plays a key role in regulating this separation of time scales. However, the detailed molecular mechanism of how the propeptide achieves this regulation is unclear. In this work, we used Molecular Dynamics (MD) simulations to characterize HP conformations and found that the HP exists in two states: one where the propeptide interacts with key residues in the HP dimerization interface and likely blocks dimerization, and one where this interface is free. We found that a propeptide mutant that dimerizes extremely slowly is essentially always in the non-dimerizable state, while the WT rapidly transitions between the two. Based on these simulations, we designed a propeptide mutant that favored the dimerizable state in MD simulations. *In vitro* assembly experiments confirmed that this mutant dimerizes significantly faster than WT. Our work thus provides unprecedented insight into how this critical step in CP assembly is regulated, with implications both for efforts to inhibit proteasome assembly and for the evolution of hierarchical assembly pathways.

## Introduction

The degradation of proteins is an essential step in signal transduction, proteostasis, and the regulation of biochemical pathways (1-3). The 26S proteasome, a massive 2.5 MDa molecular machine, is a central protease involved in intracellular protein degradation. In eukaryotes, the catalytically active 20S Core Particle (CP) is capped by two 19S Regulatory Particles (RP’s), forming the 26S proteasome. The CP consists of four heptameric rings, which are stacked coaxially in a barrel-shaped structure. The α and β subunits form the outer and inner rings, respectively, with an α_7_β_7_β_7_α_7_ stoichiometry (2, 4). The α subunits interact with the regulatory particles and help control the substrate’s entry into the barrel, while the β subunits are catalytically active and carry out proteolysis. The CP is found in all three kingdoms of life, and its overall quaternary structure is highly conserved (5, 6).

Like many molecular machines, the proteasome is not synthesized by the cell in an active form. Instead, it is assembled from a set of subunits into a functional quaternary structure. In particular, the β subunits are initially expressed in an inactive precursor form with a propeptide sequence at the N terminus, and only become catalytically active after this propeptide is autocatalytically cleaved off once the CP is fully assembled (**Fig. 1A**) (7, 8). As a result, understanding CP assembly is critical to our overall understanding of the proteasome’s function and regulation *in vivo*. The proteasome is also a well-established drug target for treating various diseases, including cancer and tuberculosis (9-11). Traditional approaches to targeting the proteasome have focused on small molecules like Bortezomib that directly bind to the active site and disrupt proteolysis (9-12). However, it has been suggested that inhibiting assembly could offer an alternative and relatively unexplored approach to pharmacologically disrupting proteasome function (8). This is particularly important for *Mycobacterium tuberculosis* (*Mtb*); it has been shown that disrupting proteasome function can ameliorate chronic *Mtb* infections, but therapies targeting proteasome assembly have yet to be developed for clinical applications (10). A better understanding of the assembly pathways and mechanisms underlying bacterial CP biogenesis could thus eventually lead to a new class of therapeutics for diseases like tuberculosis.

**Figure 1:**
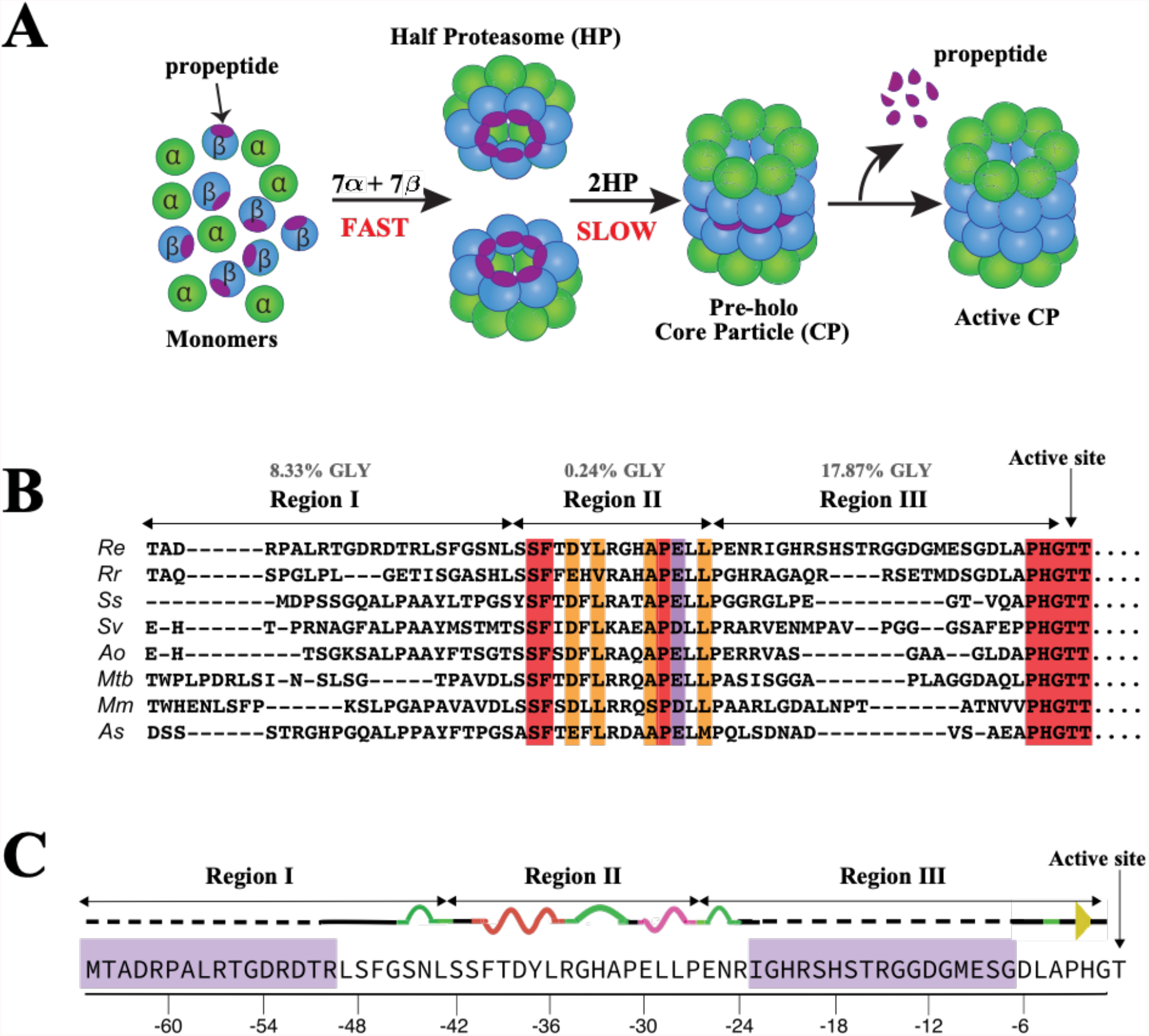
20S proteasome Core Particle assembly and propeptide sequence. (A) Schematic of proteasome Core Particle (CP) assembly. The α subunits are shown in green and β subunits in blue with the β propeptide in purple. Arrows demonstrate the steps in the progression from monomeric subunits to the active CP and, and highlight separation of time scales between Half Proteasome (HP) formation and CP assembly in bacteria (22). (B) A multiple sequence alignment of the N-terminal β subunit propeptide from 256 actinomycete bacteria. A representative subset of eight sequences is shown here: *Rhodococcus erythropolis (Re), Rhodococcus rhodnii (Rr), Saccharopolyspora shandongensis (Ss), Saccharomonospora viridis (Sv), Amycolatopsis orientalis (Ao), Mycobacterium tuberculosis (Mtb), Mycobacterium mageritense (Mm)* and *Actinopolyspora saharensis (As)*. The conserved residues are highlighted in red, residues in orange show conservation of amino acids with similar properties and the residues in purple show conservation of amino acids with weakly similar properties. The general pattern of conservation, along with available structural data, suggests the bacterial propeptide can naturally be divided into three distinct regions (22). (C) The β propeptide sequence in *Re*. Region I is made up of the -65th to the -43rd residues, Region II is from the - 42nd to the -27th residues, and Region III is from the -26th to -1st residues. The residues without electron density are highlighted in purple and are modeled for simulations.

The assembly of the CP has been studied experimentally in a wide variety of organisms (3, 13-16). In all cases, the first step in the CP assembly pathway involves the formation of Half Proteasomes (HP: α_7_β_7_) from the α and β subunits. Two HPs then dimerize to form the pre-holo-CP, after which the propeptide of the β subunit is auto-catalytically cleaved off to form the active CP (2, 7, 17, 18) (**Fig. 1A**). While the assembly pathways that form the HP differ between organisms, the HP is an obligatory intermediate in all organisms studied to date (1, 2, 14, 19, 20). Defects in proteasome assembly could lead to the accumulation of assembly intermediates within the cell or to uncontrolled proteolysis, non-specific cleavage, and aggregation of proteins if active sites are not adequately sequestered within the barrel-like structure of the CP (3, 21). As such, it is likely that the proteasome has evolved to assemble rapidly and efficiently in the cell, but many mechanistic aspects of proteasome assembly pathways remain unclear.

CP assembly kinetics have been perhaps best characterized experimentally in the bacterium *Rhodococcus erythropolis* (*Re*). Over twenty years ago, Baumeister and colleagues demonstrated that the *Re* α and β subunits remain monomeric when expressed and purified independently (7, 23). This property allows one to monitor both HP and CP formation *in vitro* as a function of time and experimental conditions(2, 7, 24). In their assembly time course experiments, Baumeister and co-workers found that the HPs are fully formed almost immediately after the subunits are mixed, with complete assembly of the HP observed at ∼30 seconds. However, fully formed CPs are visible only after a considerable time lag of about 30 minutes, and 100% CP assembly takes up to 3 hours (2, 7). This evidence suggests that HP dimerization is a rate-limiting step and demonstrates a significant separation in time scales between HP formation and dimerization to form the CP (**Fig. 1A**).

The reason for this separation of timescales is currently unknown, although existing experimental evidence supports a role for the N-terminal propeptide of the β subunit in regulating dimerization rate (7, 22). In their initial experiments, Baumeister and colleagues monitored *Re* CP assembly in three different scenarios. The first involved wild-type (WT) α and β subunits, which exhibit the separation in time scales described above. The second scenario involved WT α subunits and a mutant variant of the β-subunits with no propeptide (βΔpro), which assembled the HP at a significantly slower rate. Subsequent crystallographic studies on the *Re* CP with a β mutant where the propeptide cannot be cleaved demonstrated that the propeptide mediates critical interactions between the α and β subunits, a possible explanation as to why HP assembly is attenuated in βΔpro mutants (1, 7). In the third scenario, they added the propeptide *in trans* with α and βΔpro subunits. Surprisingly, active CP was formed significantly faster, within ∼30 seconds. The HP’s were not observed, suggesting that they dimerized so quickly that they could not be captured in native gels (7). Collectively, these findings provide evidence that propeptide regulates the dimerization rate by inhibiting HP dimerization. The molecular mechanisms through which the propeptide achieves this regulation, however, are not currently understood.

We recently published a combined computational and experimental study aimed at beginning to elucidate these molecular mechanisms (7). As part of that work, we performed a Multiple Sequence Alignment (MSA) of the β subunit from a wide range of actinomycete bacterial species (**Fig. 1B**). Based on this MSA and the available crystal structure (PDB ID=1Q5R), the bacterial propeptide can be divided into three distinct regions (**Fig. 1B)** (4, 7, 22). Region I is the most N-terminal segment of the propeptide (residues -65 to -43 in the *Re* sequence). These residues lack electron density in the crystal structure (1) and are thus likely flexible and disordered. Based on their locations in the structure, these residues likely form interactions with the α subunits. Region II (residues -42 to -27 in *Re*) is much more highly conserved than Region I, and has a well-defined, ordered structure that contacts the α and β subunits. Region III (residues -26 to -4 in *Re*) is similar to Region I, in that it shows little conservation across species and is disordered in the crystal structure. Interestingly, we found that Region III is highly enriched in glycine residues (∼17%) across all bacterial species, suggesting an evolutionarily conserved role as a disordered region of the protein (22) (**Fig. 1B**). Region III is also near the HP dimerization interface, indicating that it could play a crucial role in CP assembly.

To further test this hypothesis, we performed a series of preliminary Molecular Dynamics (MD) simulations of the HP structure to understand the dynamics of Region III and its potential role in HP dimerization (22). Upon analyzing these simulations, we found that Region III of the *Re* propeptide has a high root mean square fluctuation (RMSF) value, which is a metric that measures a residue’s flexibility averaged over time. We observed that this flexibility enables the propeptide to move closer to the HP dimerization interface and into space that could be potentially occupied by the opposing HP during dimerization. To further investigate the role of flexibility and glycine enrichment seen in the MSA, we designed several mutants targeting Region III of *Re* propeptide, primarily altering the length of Region III by making it shorter or longer. *In vitro* experiments revealed that these propeptide mutations lead to slower HP dimerization relative to WT and little to no active CP (22). These experiments provided further evidence that *Re* propeptide regulates the HP dimerization rate (2, 7, 22).

In this work, we further investigated the role of the propeptide in *Re* CP assembly by performing extensive all-atom MD simulations of the *Re* WT HP and an extremely slowly dimerizing mutant from our previous work, which we call the “SLOW” mutant (22). We used the Anton supercomputer to perform these simulations to achieve microsecond sampling for very large HP structures (25). Our simulations show specific hydrogen bonding interactions between the propeptide and several key residues at the HP dimerization interface. When forming these interactions, the propeptide physically blocks the space occupied by the incoming HP during dimerization and thus likely prevents the formation of interactions across the HP dimerization interface that are critical for HP formation and CP stability (1, 4). These interactions occurred more frequently in the SLOW mutant than in the WT. Analysis of these simulations suggested that mutating the charged residues in Region III would reduce the number and frequency of those hydrogen bonds and make dimerization faster. Hence, we designed a FAST variant by mutating two charged residues — E & D at position -9 and -12 in the β subunit — to Alanine. As anticipated, simulations of the FAST mutant HP showed fewer interactions between the propeptide, and key residues compared to WT. We then validated these predictions experimentally using *in vitro* assembly assays and found that the FAST mutant dimerizes considerably faster than WT.

These findings lead us to propose a model that suggests that the *Re* HP model exists in two conformational states: D+ (dimerizable) or D− (non-dimerizable). In the D− state, interactions between the propeptide and key dimerization residues block dimerization, while in the D+ state those residues are free to interact with another HP. In solution, the D+ and D− states exist in equilibrium, and mutations or other perturbations can influence this equilibrium. Our results show that, in WT, the D+ and D− are both present and interconvert on μs time scales, but in the SLOW mutant, the D− conformation dominates and essentially does not interconvert with the D− on the time scale of our simulations. As anticipated, the FAST mutant more frequently resides in the D+ conformation. This model explains our previous experimental results and provides unprecedented mechanistic insights into how HP dimerization is regulated (22).

As mentioned above, Region III of the propeptide has an enriched level of glycine in the majority of bacterial species, and many species, including *Mtb*, have charged or polar residues in roughly the same region of the propeptide as in *Re* (**Fig. 1B**). This suggests that similar mechanisms are at play in the majority of bacterial species. If the *Mtb* CP uses a similar mechanism, it might be possible to discover small molecules that bind selectively to the D− conformation and stabilize it, thus slowing or preventing dimerization and inhibiting proteasome function. This could lead to an entirely new class of proteasome inhibitors that target bacterial infections. The propeptides of the β subunits in archaea and eukaryotes have very divergent lengths and do not necessarily share the Region I/II/III architecture with bacteria. Future computational and experimental work will be necessary to elucidate the extent to which disordered regions of the propeptide play a role in regulating HP dimerization in other species of bacteria and in archaea and eukaryotes.

## Materials and Methods

### Half Proteasome structure for Molecular Dynamics simulations

The X-ray crystal structure of the *Rhodococcus erythropolis* Core Particle (CP) was used as a starting structure and is available in PDB 1Q5R (1). The starting structure for the Half Proteasome (HP) simulations was obtained by two of the four rings from the CP coordinates (one α ring and one β ring, of course bound to each other), with all 14 chains. Additionally, 1Q5R has a point mutation in the catalytic active site triad, which was essential for crystallizing the *Re CP* with the propeptide (1). In our simulations this residue, i.e., ALA-33, has been mutated back to the WT sequence (LYS-33).

### Modeling the missing propeptide residues

In the starting structures for these simulations, each propeptide of each β subunit has 33 out of 65 residues are not present (**Fig. 1C** purple background) due to missing electron density in the x-ray crystal structure. To model these residues, we start with a single α-β dimer from the 1Q5R structure (specifically, chains A and H). Next, the 33 missing residues were modeled using the default parameters of the comparative modeling method implemented on the Robetta server (27, 28). Robetta generates multiple models, so we chose the model with the lowest error (Å) for the next step. We performed a set of symmetry operations in Pymol (26) to replace the *Re* α-β dimers in 1Q5R with the Robetta modeled α-β dimer. The resulting structure was investigated for ring penetration, knots, and any steric clashes that can cause potential artifacts. To do this, we used Anton2 validation checks using a built-in DMS module on Anton2 (25).

We used the WT modeled structure of the HP as a template for the SLOW mutant and extracted an α-β dimer from it. To model the additional residues of the SLOW mutant, we specified the residues in the FASTA sequence, which served as the input for an additional set of comparative modeling runs in Robetta (27, 28). This resulted in a Robetta model for the SLOW mutant α-β dimer. The SLOW mutant HP models were generated using the same procedure as the WT models and subjected to the same validation steps for obtaining the equilibration and production simulation runs. The FAST mutant models were generated directly from the WT models by mutating the corresponding residues (glutamate E-9 and aspartate D-12 to alanine) in the CHARMM-GUI solution builder module (29) (see below).

### Molecular Dynamics simulation setup

The simulation inputs were generated using the CHARMM-GUI solution builder module (29-31). All the systems were neutralized with 100mM NaCl (which is the salt concentration used in the experiments) and 15Å water on each side of the protein for a rectangular box. The detailed components are provided in the supplementary material (**S.10**). First, we performed a short minimization of 5000 cycles, switching from steepest descent to conjugate gradient after 2500 cycles. For minimization, the system was held at constant volume, without any restraints on the atoms, and nonbonded cutoff of 10Å. Next, we performed 5ns of NVT and 60ns NPT equilibration with a 2fs time step using the GPU version of AMBER18 (32, 33) and the CHARMM forcefield 36m (30). Each simulation took approximately 48 hours to finish 65ns of equilibration on an NVIDIA RTX 2080Ti GPU. The detailed parameters for the equilibration are given in the supplementary material. After the equilibration in AMBER, the coordinates and restart files were used to initiate 2.5 µs Molecular Dynamics simulations on Anton2 (25). The NPT ensemble, CHARMM 36m forcefield and TIP3P water model were used for the Anton simulations. The pressure and temperature were kept constant at 1bar and 303.15 K using Multigrator integrator and default NPT parameters (34). The RESPA (REference Systems Propagator Algorithm) integration method was employed with a timestep of 2.5 fs, and the coordinates were saved every 0.24ns. All systems were simulated under periodic boundary conditions in the NPT ensemble. Additional details of the equilibration and MD simulations are provided in section S.10 of the supplementary material. All simulations for the WT and mutants had a backbone RMSD around 3.5Å – 6.5Å (**Fig. S4**) and required about 500 nanoseconds to converge (**Fig. S1**). As a result we did not include the first 500 nanoseconds of the runs in our quantitative analyses.

### Statistical analysis

We used a categorical regression in R to estimate the significance of the differences in dimerization states between the WT and mutant simulations (**Tables S.9**). These tests were used to determine if the behavior of WT is statistically different from SLOW and FAST, both in terms of average properties and in terms of systematic changes over simulation time. MD simulation data is time-dependent and autocorrelated, and as such, it is essential to employ statistical tests that are robust to these factors. We thus employed categorical linear regressions to perform our analyses and used the Newey-West estimator, which is robust to both heteroscedasticity and autocorrelation, to efficiently estimate the standard errors and compute the p-values (35-38). All linear regression analyses, estimates, and tests were done using the lm function in R (39). Further details on this analysis, and the specific scripts, are available in the supplementary material section S.9.

### Expression and purification of the *Rhodococcus erythropolis* α1 subunit

pT7 plasmid containing the α1 gene of *Rhodococcus erythropolis* was obtained as a kind gift from Wolfgang Baumeister. The α1 gene was then subcloned in a pTBSG plasmid background with an N-terminal TEV-cleavable 6xHis-tag by Philip Gao and Anne Cooper at the Protein Production core lab, University of Kansas. Plasmids were freshly transformed in *E. coli* BL21 (DE3)-pLyse S (NEB, Catalogue # C3010I) cells. Cells were grown in 4 Liters Luria Barteni (LB) media containing 100 μg/ml Carbenicillin and 30μg/ml Chloramohenicol. Cells were grown in a shaking incubator at 200rpm and 37°C, induced with 1 mM isopropyl-β-D-thiogalactopyranoside (IPTG) at OD600 ∼ 0.6 and were allowed to grow overnight at 15°C. Cells were harvested by centrifugation at 4000 rpm for 15 mins using a (JS-4.750 rotor in an Avanti-J15R centrifuge), resuspended in lysis buffer (50mM Tris, 100mM NaCl, 10mM MgCl_2_, 1mM ATP, 5mM BME, 30% glycerol pH7.05 at RT), and sonicated at 45% amplitude with 2s on - 10s off cycles for 4 mins using a Fischer Scientific Sonic dismembrator model 500. Cellular debris were removed by centrifugation at 10,000 rpm for 30 min in a JA-10.100 rotor in an Avanti-J15R centrifuge. Clarified lysate was passed through a 10 ml Ni^2+^-Sepharose 6 Fast flow resin (GE Healthcare Catalogue number 17-5318-02), washed with 50 ml of binding buffer (50mM Tris, 100mM NaCl, 10mM MgCl_2_, 1mMATP, 5mM BME, 30% glycerol pH7.05 at RT) followed by a second wash with 50ml wash buffer (50mM Tris, 100mM NaCl, 10mM MgCl_2_, 1mMATP, 5mM BME, 30% glycerol, 10mM Imidazole pH7.05 at RT) and eluted with a total of 50 ml elution buffer (50mM Tris, 100mM NaCl, 10mM MgCl_2_, 1mMATP, 5mMBME, 30% glycerol, 1M Imidazole pH7.05 at room temperature). Fractions containing purified protein were incubated with 1:20 v/v of 62μM TEV protease (1ml TEV/20mL protein) and dialyzed overnight in a binding buffer. Dialyzed protein was subjected to the exact same Ni^2+^-affinity chromatography to separate the protein from cleaved His-tags and TEV protease. Fractions containing the protein were pooled together and mixed with 20% v/v glycerol. Aliquots of 1ml were stored at -80°C for later use.

### Expression and purification of the *Rhodococcus erythropolis* β1 subunit

DNA corresponding to the β1 gene of *Rhodococcus erythropolis* optimized for expression in *E*.*coli* was purchased from Genscript. The optimized gene was PCR amplified and subcloned into the NdeI/XhoI sites of pET-22b (a generous gift of Roberto De Guzman, University of Kansas) to introduce a C-terminal 6xHis-tag for protein purification and generate the pAKRE-β plasmid. EtoA and DtoA mutations at positions -12 and -9 respectively were generated using site directed mutagenesis to generate pJR 868 by Jeroen Roelofs at University of Kansas Medical Center. Plasmids were freshly transformed in *E. coli* BL21 (DE3) and cells were grown in 4L LB-media containing 100 μg/ml carbenicillin. Cells were grown in a shaking incubator at 200rpm and 37°C, induced with 1mM isopropyl-β-D-thiogalactopyranoside (IPTG) at OD600 ∼ 0.6 and cell growth was continued for 4 hours at 30°C. Cells were harvested by centrifugation at 4000 rpm for 15 mins using a (JS-4.750 rotor in an Avanti-J15R centrifuge) and pellets were resuspended in 50 ml binding buffer (25mM Na2PO4, 300mM NaCl, 5mM Imidazole pH7.4). Cells were lysed by sonication at 45% amplitude with 2s on-10s off cycles for 4 mins using a Fischer Scientific Sonic dismembrator model 500. Cellular debris were removed by centrifugation at 10,000 rpm for 30 min in a JA-10.100 rotor in an Avanti-J15R centrifuge. Following centrifugation, the protein was purified using a GE AKTA pure FPLC system. The cleared lysate was loaded on a Nickel affinity column (GE His-trap 5ml column), the column was then washed with 10 column volumes (CV) of binding buffer (25mM Na_2_PO_4_, 300mM NaCl, 5mM Imidazole pH7.4). The bound protein was then eluted with a linear gradient of the elution buffer (25mM Na2PO4, 300mM NaCl, 500mM Imidazole pH7.4) over 20 CV. Fractions containing the protein were dialyzed in anion exchange binding buffer (20mM Tris-HCL, 20mM NaCl, pH 8.0 at 4^0^C) overnight at 4^0^C. On the second day, dialyzed protein was cleaned up using a GE 5ml Hi-trap anion exchange column. Unbound protein and contaminants were washed with 10 CV of binding buffer (20mM Tris-HCL, 20mM NaCl, pH 8.0 at 4^0^C). Bound protein was eluted using a linear gradient of the elution buffer (20mM Tris-HCL, 1M NaCl, pH 8.0 at 4^0^C) over 20 CV. Peak fractions were analyzed using SDS-PAGE. Fractions containing the protein were pooled together. 20% V/V glycerol was added to the pooled fractions and aliquots of 300µL were frozen in liquid nitrogen and stored at -80^0^C for later use.

### *In vitro* reconstitution reactions

The α subunit was mixed with the wild type β or FAST mutant β separately in equimolar ratios to obtain a final subunit concentration of 4μM. Assembly reactions were allowed to proceed for 3hrs at 30°C. At the end of the 3hrs time course, equal volume of loading dye (20mM HEPES, 0.1% Bromophenol Blue, 20% Glycerol) was added to the reactions. Samples were loaded on a 4-20% Tris Glycine native gel (Invitrogen). Gels were run at 4°C, 120V for 12 hours, stained with Sypro Ruby (Thermofisher Scientific, catalogue number S12000) protein stain as described by the manufacturer, visualized using Biorad ChemiDoc imager and quantified using ImageLab software.

## Results

### Interactions between the propeptide and key dimerization residues

Shorter simulations from our previous work revealed that Region III of the WT *Re* propeptide is highly flexible, lies near the HP dimerization interface, and is generally found protruding outside the HP barrel (23). In this work, we used the Anton2 resource (26) to perform 2.5μs Molecular Dynamics (MD) simulations for three independent replicates of the WT HP structure. Visual inspection of the resulting trajectories revealed that the β propeptides of some subunits spent considerable time near the HP dimerization interface during the simulation (**Fig. S1**). Previous analysis of the *Re* crystal structure revealed a set of “key residues” in the dimerization interface that mediate interactions between a given β subunit and the opposing β’ subunits of the other HP (4) (**Fig. 2A and B**). These interactions include hydrogen bonds, salt bridges, and hydrophobic interactions between the β rings (4). Witt *et al*. demonstrated that perturbing any of these critical interactions by mutating the key residues has a destabilizing effect and, in most cases, completely prevents CP formation, suggesting that they are critical to the HP dimerization process (4).

**Figure 2:**
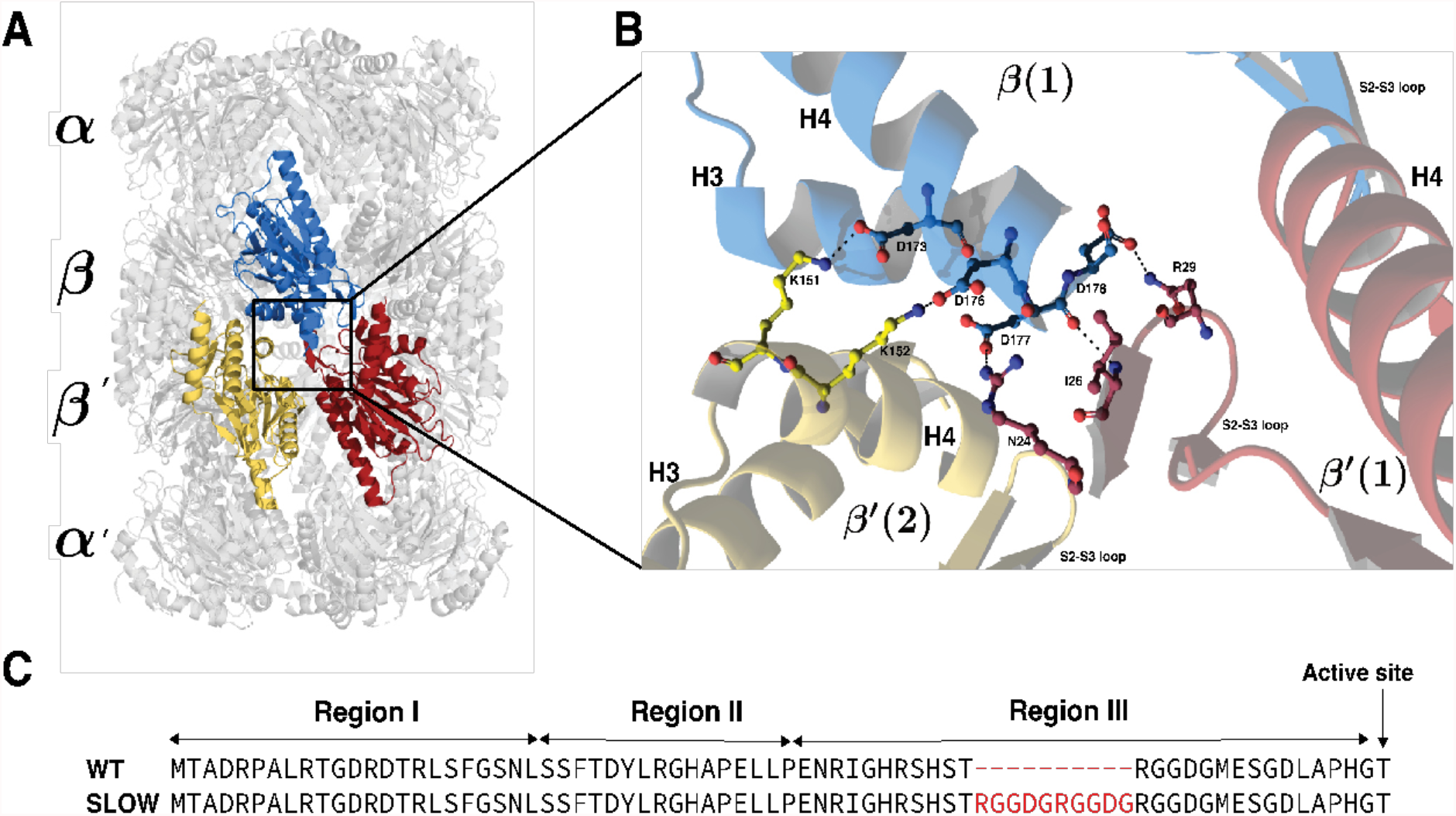
*Re* CP structure and key residues. (A) Side view of the WT *Rhodococcus erythropolis* (PDB entry:1Q5R) Core Particle (CP). The colored β subunits (blue, yellow and red) highlight interactions between β and β’ subunits which occur at the Half Proteasome (HP) dimerization interface. (B) This inset shows the zoom in view of the key residues associated with β-β’ interactions that are part of S2-S3 loop and H3-H4 helices (1) (C) *Re* β propeptide sequence for Wild Type (WT) and the SLOW mutant, which dimerizes at a much slower rate.

On analyzing our WT simulation results, we found that a stretch of about 7-10 residues of Region III is more mobile flexible than Regions I or II (**Fig. S2**) and formed hydrogen bonds with the key residues in the HP dimerization interface (**Fig. 2A** and **B**). Specifically, we observed that the ^-13^GDGMESG^-7^ motif in Region III of the WT propeptide frequently interacts with the key residues (**Fig. 2C**). In our previous work, we had designed an “Extended Loop 1” (EL1) mutant with an extended Region III by repeating a charged and glycine-rich portion of Region III twice (^-15^RGGDG^-11^ in the WT sequence) (**Fig. 2C**). Interestingly, *in vitro* assembly assays showed HP formation in this mutant, but no CP was observed even after prolonged incubation (22). In this work, we refer to this EL1 variant as the “SLOW” variant because HP dimerization occurs very slowly for this mutant (if at all) (**Fig. 2C**).

We noted that the duplicated motif in the SLOW mutant overlapped significantly with the propeptide region that was observed to be interacting with the key residues in our simulations. We thus performed an additional set of three 2.5 µs simulations for this mutant. Interestingly, on visual inspection of the resulting trajectories, we observed that the propeptide of the SLOW mutant interacted with the key residues in the dimerization interface much more frequently than in WT. This led us to the hypothesis that the HP exists in two distinct conformational states: [1] an “interacting” state where the propeptide forms hydrogen bonds with the key residues and [2] a “non-interacting” state where the propeptide does not form hydrogen bonds with the key residues. In the interacting state, the key residues are occupied by the propeptide, which physically blocks the opposing HP from forming interactions with those residues. Since mutations to the key residues block dimerization (4), we thus hypothesized that this state cannot dimerize, and thus refer to this state as non-dimerizable (D−). On the other hand, if no hydrogen bonds are formed between the propeptide and the key residues, the HP is free to interact with another HP. We thus call this the dimerizable (D+) state. **Fig. 3A and B** illustrate the D+ and D− states.

**Figure 3:**
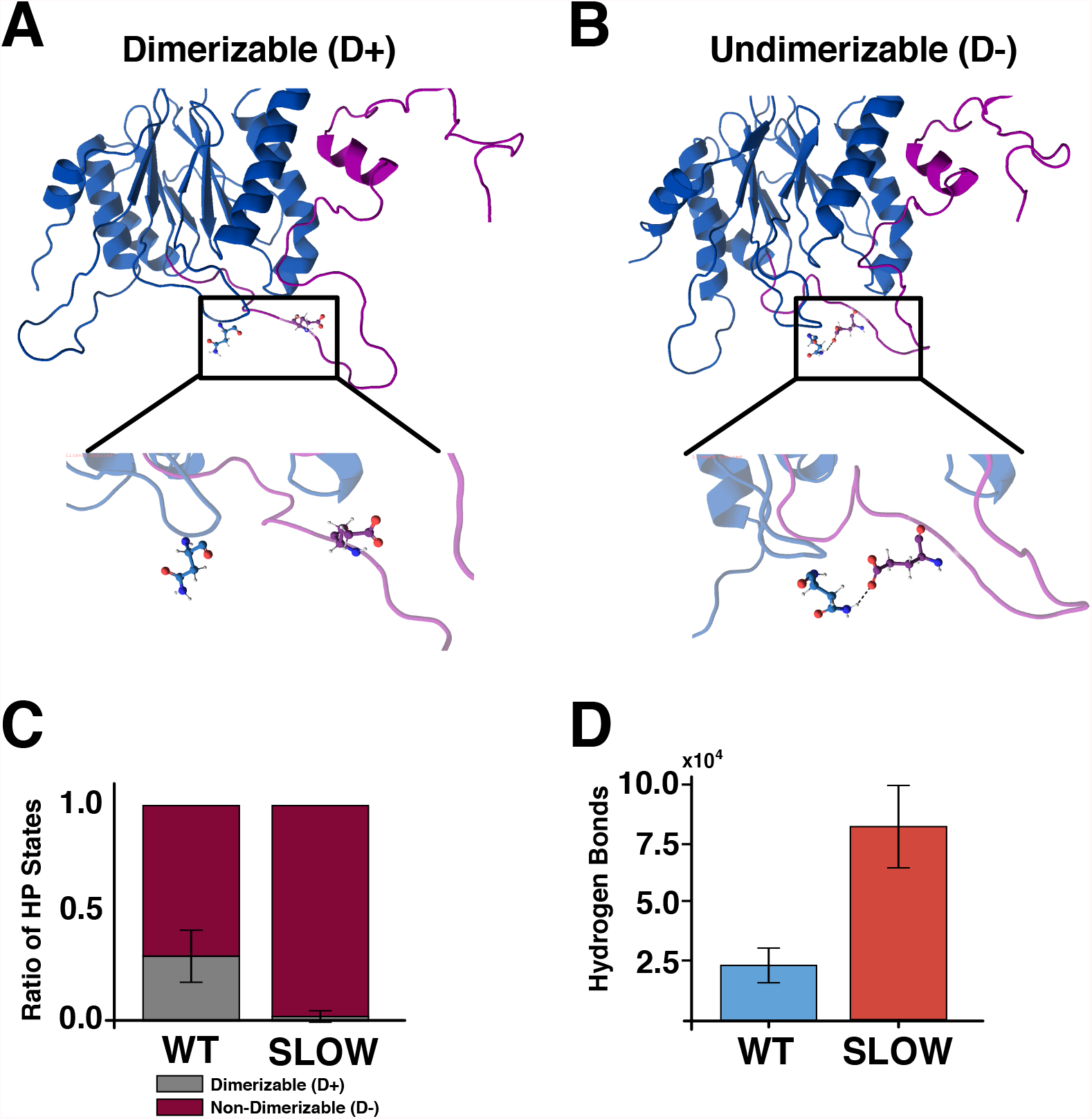
Proteasome conformational states during assembly. (A-B) Ribbon diagrams of the β subunit (blue) with a full length propeptide (purple). Key residues are shown as colored atoms. (Left) β subunit of a HP in a non-dimerizable state with inset highlighting interactions. (Right) β subunit of HP in a dimerizable state for comparison. (C) Bar graph of percent of simulated time frames in D+ and D−states for both WT and SLOW β subunit mutants. Error bars signify SEM for 3 MD simulations of 2.5 µs. (D) Bar graph showing the total number of hydrogen bonds formed between the propeptide and the key residues as an average over 3 MD simulations for both WT and SLOW Half-Proteasomes (HPs). Error bars show the Standard Error of the Mean for 3 MD simulations of 2.5 µs.

If this hypothesis is correct, we should see a significant difference in the frequency and character of the interactions between the propeptide and the key residues when comparing the WT and SLOW variants. We quantified these interactions by calculating the number of simulation frames (corresponding to every 0.24 ns of simulation time) in which a hydrogen bond (based on a 2.4Å distance cutoff) between a key residue and a propeptide residue is formed. If there are hydrogen bonds between any key residue and a propeptide residue, we consider that conformation to be in the D− state. If there are no such bonds, we consider it to be in the D+ state. Consistent with our hypothesis, simulation results revealed that the SLOW mutant propeptide resides in the D− much more frequently than the WT (**Fig 3C**). We found that the simulation systems required around 500 ns to equilibrate, so frames from the first 500 ns are ignored in this analysis (**Figs. S3** and **S4**). In particular, the WT HP exists in the D+ state about 25% of the time across the three simulated replicates, and the D− state 75% of the time (**Fig. 3C**). In contrast, the HP of the SLOW mutant nearly always (>99% of simulated time) resides in a D− state (**Fig. 3C**). To further characterize the difference between the two mutants, we also counted the total number of hydrogen bonds made between any propeptide residue and any key residue. We found that the SLOW mutant forms more than 3 times the number of hydrogen bonds as the WT. Our observations for both the frequency of D− states and the number of hydrogen bonds formed were consistent across all three replicates (**Fig. 3C** and **D**).

Note that our analysis in **Fig. 3** focuses entirely on hydrogen bonds, but of course other forms of interaction could also occur between the propeptide and the key residues. We found, however, that whenever the propeptide was close enough to the dimerization interface to interact, hydrogen bonds were formed between charged residues at the dimerization interface and residues or backbone atoms of the propeptide. So, while the analysis above focuses on hydrogen bonds, the results are representative of essentially any interaction between the propeptide and the key residues. Also, our definition of the D− state is rather stringent: if any propeptide residue forms an interaction with any of the key residues on any of the seven β subunits, we consider that to be a non-dimerizable state. As **Fig. 3D** demonstrates, the SLOW mutant simply has more interactions overall. So alternative definitions (requiring more total interactions and/or more subunits with interactions to qualify for the D− state) would not qualitatively change the large difference in the propensity of interactions we observed. These results thus suggest that the large difference in dimerization rate between the WT and SLOW mutant observed experimentally are because the longer propeptide in the SLOW mutant (with many more charged residues) interacts much more frequently with the key residues and blocks dimerization (22) (**Fig. 2C**). These findings also suggest a straightforward explanation for the separation of time scales in WT CP assembly: the HP spends most of its time in the D− state, and as such there are relatively few HPs competent to dimerize with one another at any given time. This would naturally lead to a situation where HP dimerization is much slower than HP assembly (2, 7, 22).

### Mutating charged residues on the propeptide generates a fast-dimerizing variant

One clear prediction of the D+/D− hypothesis described above is that mutations that reduce the level of interactions between the propeptide and the key residues should favor the D+ state and thus lead to faster dimerization. To test this idea directly, we computationally mutated two charged residues in WT Region III, glutamate E-9 and aspartate D-12 to alanine using CHARMM GUI (30) to generate what we term the FAST mutant (**Fig. 4A**). In the WT HP simulations, both residues interact with the key residues, specifically by forming side-chain hydrogen bonds with several amino acids that form salt bridges across the HP dimerization interface, particularly R29 and N24 of the β subunit. We thus hypothesized that mutating these residues to alanine could potentially contribute to faster dimerization. We performed three replicate MD simulations of the FAST mutant HPs for 2.5 µs. Analysis of these simulations revealed that interactions between the propeptide and the key dimerization residues were less frequent compared to WT, with the FAST mutant spending close to 50% of the simulation in the D+ state (**Fig. 4B**). Furthermore, the FAST HP formed fewer hydrogen bonds with key residues compared to the WT HP (**Fig. 4C**). As expected, interactions between the -9 and -12 residues of the propeptide and the key residues were much less frequent in FAST than WT (Fig. S7). This suggested that the FAST mutant would indeed dimerize faster than WT.

**Figure 4:**
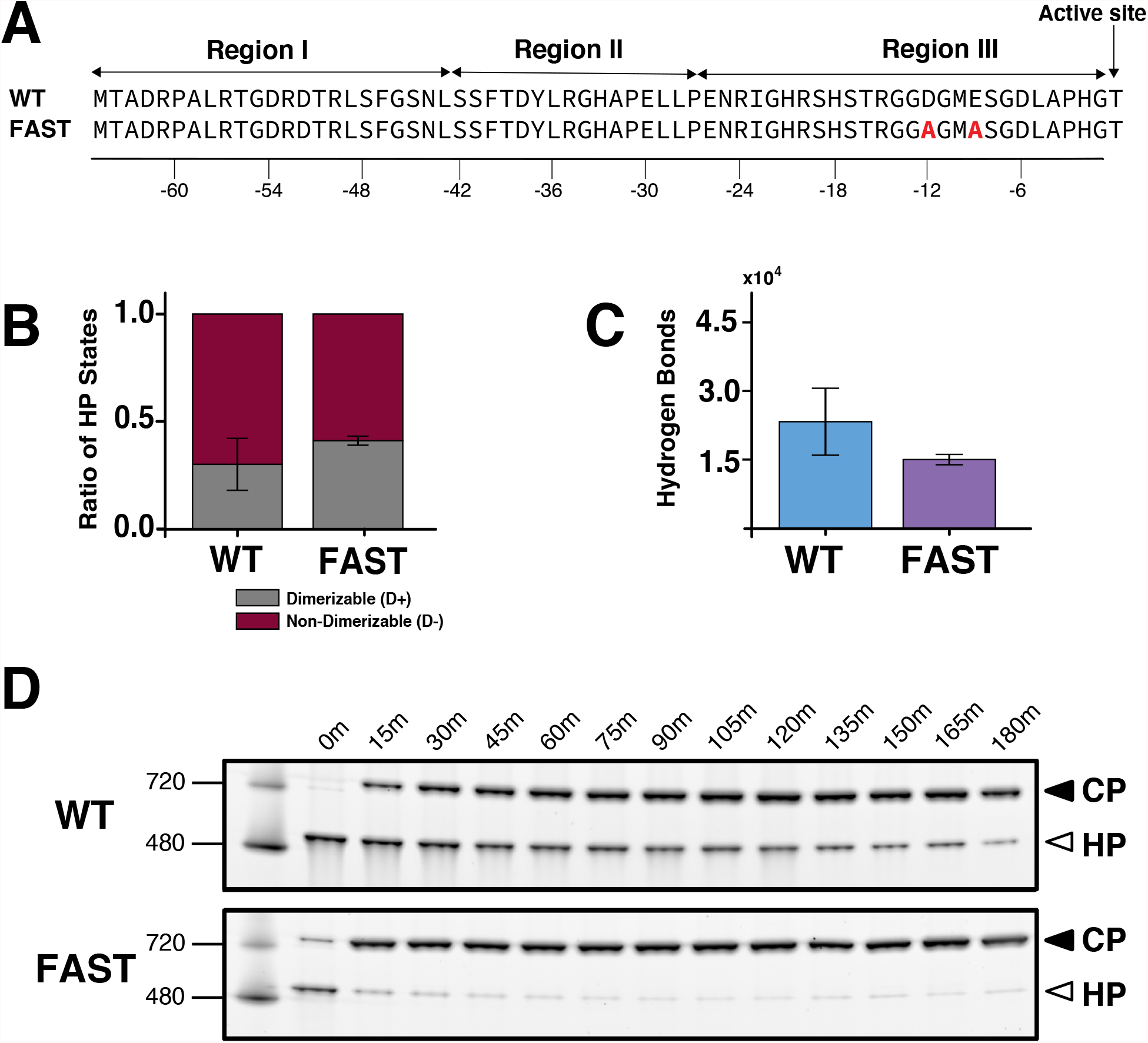
Proteasome conformational states during FAST assembly. (A) Sequence of β subunit propeptide regions in WT and FAST mutants. Residues in red are the mutated E (glutamate), D (aspartate) to A(alanine). (B) Bar graph of percent of simulated time frame in D- and D+ states for both WT and FAST β subunit mutants Half-Proteasomes (HPs). Error bars signify SEM for 3 MD simulations of 2.5 us. (C) Bar graph showing the total number of hydrogen bonds formed between the propeptide and the key residues as an average over 3 MD simulations for both WT and FAST HPs. Error bars show SEM for 3 MD simulations of 2.5 us. (D) 4-20% Tris-Glycine native gels from *in vitro* assembly assays at increasing time points (time points labeled above each lane in minutes) for WT and FAST β subunit mutants. Gels were stained with Spyro Ruby protein and visualized with a BioRad Imager.

To test our predictions experimentally, we generated a double point mutation D-12A/E-9A variant (**Fig. 4A**). To compare the assembly rates of WT and FAST mutants, α and β mutant substrates were mixed *in vitro* and incubated at 30^0^C at increasing time points from 0 minutes to 180 minutes and performed Native PAGE on the reaction products (3, 8, 23). This assay provides a readout of assembly kinetics of both the WT and FAST mutant HP and CP (**Fig. 4D**). Consistent with previous experimental results (4, 7, 22) we observed that WT HP formation is essentially complete at our earliest time point (“0” minutes), which is obtained by mixing the subunits, immediately loading the reaction on the gel, and then running the gel. In the WT case, CP starts to appear after 15 minutes, and continues to increase in concentration as indicated by the increase in band intensity (**Fig. 4D**). In contrast, the FAST mutant shows significantly more CP formation at the 0th and 15-minute time points, with CP assembly essentially complete by 30 minutes. These findings demonstrate that the FAST mutant does indeed dimerize considerably more rapidly than the WT.

### Dynamics of interactions between the propeptide and the key residues

To further characterize the observed differences between the β subunit variants discussed above, we analyzed the dynamics of hydrogen bond formation. Specifically, we looked at the total number of hydrogen bonds formed as a function of simulation time; representative examples for each variant are shown in (**Fig. 5 A, B**, and **C**). In each case, the colored lines represent the actual number of hydrogen bonds observed in each simulation frame, and the black line represents a Locally Weighted Scatterplot Smoothing (LOWESS) estimate of the moving average (40). For the WT simulations, we see that the propeptide residues form around 4-10 total hydrogen bonds with the key residues by about 500 ns of simulation time (as mentioned above, theses first 500 ns of the simulations are taken to be equilibration time and are not included in our analyses). After the initial equilibration, the WT system spends most of the time in this D− state, only visiting a state with 0 hydrogen bonds transiently (**Fig. 5A**). The SLOW mutant rapidly forms 12-20 hydrogen bonds, and on average has about twice as many interactions as the WT (**Fig. 5B**). As a result of this greater number of interactions, the D− state is much more stable in the SLOW case, and we do not observe any transitions to the D+ state once these interactions form (**Fig. 5B**). Finally, in the case of the FAST mutant, we see fewer hydrogen bonds on average (0-4 bonds) and very frequent transitions between the D− and D+ states (**Fig. 5C**).

**Figure 5:**
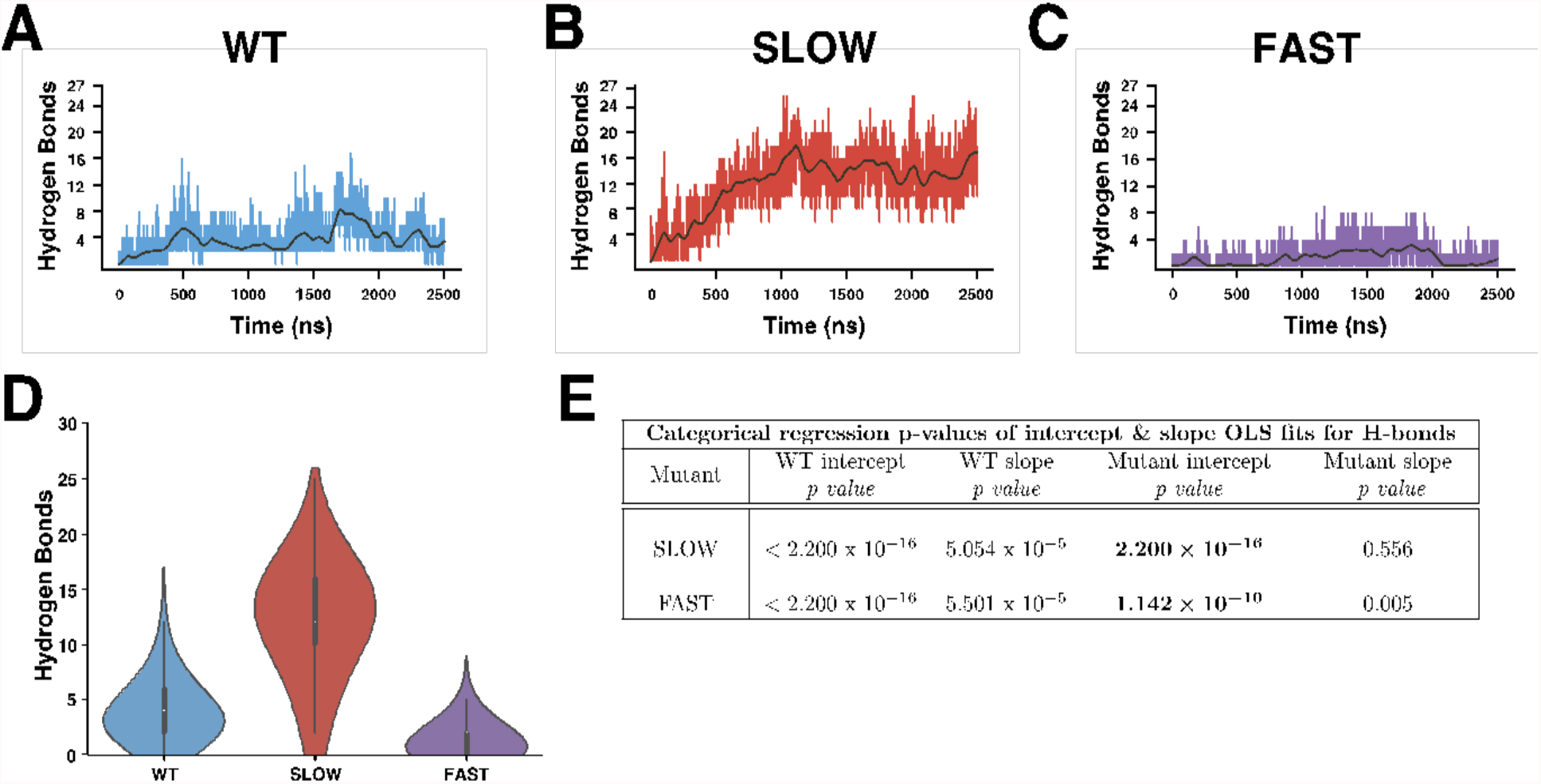
Hydrogen bond dynamics in MD simulations, shown for one replicate of each type. (A-C) LOWESS plots of the number of hydrogen bonds formed by the HP over simulated time (in ns) for a single MD simulation of WT (left), SLOW (middle) and FAST (right) β subunit mutants. The bold line represents a non-parametric LOWESS fit. Similar plots for remaining replicates are included in Supplementary Material. (D) Violin plot of distribution of the total number of hydrogen bonds formed over each of three independent MD simulations of WT (blue), SLOW (red) and FAST (purple) HPs. The white dot in the violin plots represents average number of hydrogen bonds for that computational replicate. E) Categorical regression p-values to compare the WT and the SLOW, FAST mutants shown in (D).

Taken together, our results suggest a strong statistical distinction between hydrogen bonding propensities between these three variants (**Figs. 3 C** and **D, 4 B** and **C**, and **5 A-C**). Indeed, if we collect all the frames from all three replicates and plot the distributions of the number of hydrogen bonds formed, these distributions look quite distinct for the WT compared to the SLOW and the FAST (**Fig. 5D**). However, testing the statistical significance of this difference is non-trivial due to the autocorrelated nature of MD simulations. Most straightforward statistical tests for example to test the difference of means between two distributions (e.g., a t-test or a Wilcoxon rank-sum test) assume that the constituent samples that form the observed distribution are independent and identically distributed. This is not the case for MD data; if the number of hydrogen bonds present in some simulation frame is 10, it is improbable that the number of hydrogen bonds will fall to 0 in the next frame 0.24 ns later.

We thus employed a robust regression-based statistical approach that specifically controls for the autocorrelated nature of time series like those obtained from MD simulations to overcome this problem. This technique is based on standard linear regression but is modified to compare a pair of replicates. The approach is based on the Newey-West estimator, which is robust to heteroscedasticity (i.e., differences in the standard deviation across time) and autocorrelations. Further details on the regression models themselves can be found in the supplementary material. This approach provides four different p-values corresponding to different statistical questions. The first two involve whether the intercept and slope of the WT replicate are statistically distinct from 0; p-values for one of the WT replicates are shown in **Fig. 5E**. The more interesting question involves comparing the mutant replicates to the WT. In this case, the p-value reports how unlikely it is for the WT and mutant coefficients to be the same. In other words, this approach indicates whether the time-dependent pattern of hydrogen bonding is different between the replicates.

A table reporting the p-values for a comparison between one of the WT replicates and one each of the SLOW and FAST mutant replicates is shown in **Fig. 5E**. For these comparisons, we see that the *intercept* for the SLOW and FAST mutant have a low probability of being identical to the intercept for the mutant (<10^−10^ in both cases). Although the slopes are not significantly different, this finding suggests that the average hydrogen bonding behavior of the mutant replicates is unlikely to be the same as WT. Although **Fig. 5E** reports the results for comparing only one set of replicates, we have performed all possible pairwise comparisons between replicates (see the supplementary information). After correcting for multiple hypothesis testing, we see significant differences between the WT and the two mutants in every case. This persists across a number of alternative categorical regression models that we have considered, suggesting that the differences between the WT and the mutants observed in **Fig. 5D** are not likely to be a statistical artifact but rather represent fundamental differences in the dynamics between the WT and mutant systems.

The fact that the proteasome is a multimeric complex allows for the possibility that individual subunits might show coupled transitions between the D+ and D− conformations. In other words, if one β subunit propeptide interacts with the key residues, does that influence the other β subunits to interact as well? To investigate this, we made a simple kymograph plotting the number of hydrogen bonds between the propeptide and the key residues in each β subunit as a function of time. **Fig. 6** shows representative results of the WT (**Fig. 6A**), FAST (**Fig. 6B**), and SLOW (**Fig. 6C**) HP simulations. As expected, the SLOW mutant had more hydrogen bonds across the subunits than WT and FAST. We observed no evidence of cooperativity between the subunits in any of our replicates (see the supplementary info for kymographs of the other two replicates for each system). Interestingly, we observed a heterogeneous distribution of hydrogen bonds among the seven β subunits for each variant. For instance, for the WT replicate in **Fig. 6A**, the β1, β2, and β6 subunits show high numbers of hydrogen bonds, while the remaining β subunits have very few. We observed this heterogeneity even for the SLOW mutant; in **Fig. 6B**, we see almost no hydrogen bonding for β7, even though most subunits are in the D− state. The variants also showed significant differences in the time scales of interactions between the propeptide and the key residues. In WT, the D− state persists for hundreds of ns, while in the SLOW case, once a subunit enters the D− conformation, it tends to stay there for the duration of the simulation. This indicates that transitions between D+ and D− are fairly frequent for individual subunits in WT and much less frequent in the SLOW variant. On the other hand, D− conformation is considerably more transient in the FAST case, generally persisting for only tens of ns (**Fig. 6C**). Taken together, these results suggest that individual subunits independently sample these two conformations and that mutations that influence dimerization rates have a significant influence on the stability and transition rates between these states.

**Figure 6:**
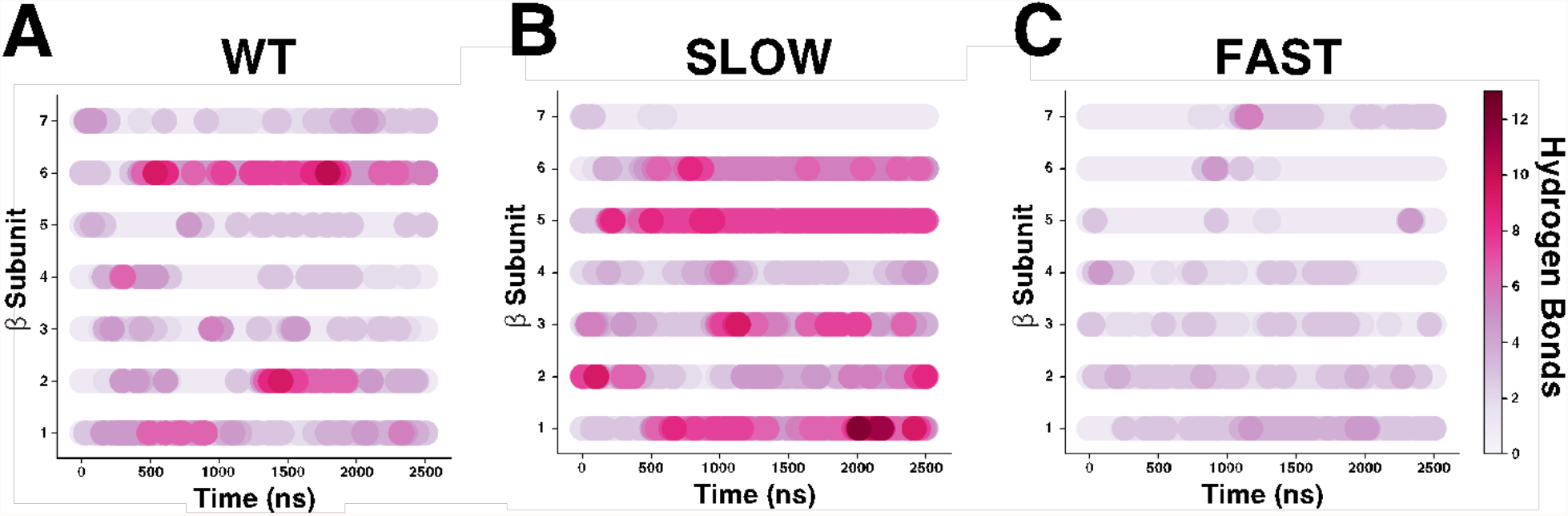
Hydrogen bond dynamics in each β subunit. Kymographs of the number of hydrogen bonds formed over simulated time for each β subunit in one replicate of the HP of WT (A), SLOW (B), and FAST (C) mutants. Colors correspond to the number of hydrogen bonds formed, with the brighter red representing more bonds.

## Discussion

By virtue of its role as a major protein degradation machine, the proteasome is involved in nearly every cellular process, including regulating the cell cycle, protein homeostasis, and cell signaling. The assembly of the proteasome Core Particle (CP) has been studied in several species (7, 8, 21). However, while several aspects of the CP assembly process have been experimentally well established, very little work has been done to understand the regulation of CP assembly at an atomistic level. This work is, to our knowledge, the first attempt at using MD simulations to study an aspect of the CP assembly process in at this level of structural resolution.

A critical step in CP assembly in all organisms is the dimerization of two Half Proteasomes (HPs). In the bacterial species *Rhodococcus erythropolis* (*Re*), a classic set of *in vitro* assembly assays showed that the HP is fully assembled quickly, while CP assembly takes considerably longer (7). This and subsequent work demonstrated that the β propeptide plays a critical role in regulating the rate of HP dimerization and other steps in the assembly process (22). Although this was shown experimentally almost 25 years ago, the molecular mechanism underlying this separation of time scales, and the specific aspects of the propeptide that allow it to regulate dimerization rate, have remained unclear.

Here, we performed all-atom Molecular Dynamics (MD) simulations of the *Re* HP, focusing on WT and β propeptide mutants. Specifically, we focused on a variant of the β propeptide in which a flexible, charged “RGGDG” motif present in the WT propeptide was repeated several times (**Fig. 2C**). Experimentally, we had found that this mutant, which we call the SLOW mutant, shows no dimerization on the time scale of days (22). In our simulations of the WT sequence, we observed that the propeptide frequently makes interactions with several of the canonical “key residues” in the HP dimerization interface that have been experimentally shown to be critical for CP formation (**Fig. 2A**). This led us to hypothesize that the HP exists in two states, a D− state where the propeptide interacts with the key residues and dimerization is blocked, and a D+ state where the propeptide is not interacting with the key residues and dimerization is possible (**Fig. 3A**). In our simulations of the WT HP, we found that it resides in the D− state about 75% of the time. Interestingly, the SLOW mutant resides in this state nearly 100% of the time. In addition, the sheer number of interactions between the propeptide and the key residues was much higher in the SLOW mutant than in the WT (**Fig. 3**). This led us to hypothesize that the equilibrium between the D− and D+ states is likely to lead to the separation of timescales in WT CP assembly and that, by shifting the equilibrium almost entirely to the D− state, dimerization is completely abrogated in the SLOW mutant.

To test this hypothesis directly, we designed another propeptide mutant that we expected would reduce the number of interactions between the propeptide and the key residues, thus shifting the equilibrium to the D+ state and increasing the dimerization rate. Specifically, we chose two negatively charged residues in the propeptide that frequently interacted with the key residues in our WT simulations: E-9 and D-12 (**Fig. 4A**). Next, we computationally mutated these residues to alanine to generate what we call the FAST mutant and, as expected, we saw a shift in the equilibrium that favored the D+ state and a corresponding decrease in the total number of interactions between the key residues and the propeptide in our MD simulations (**Fig. 4B, C)**. We then made this mutant in the lab and tested its assembly kinetics experimentally and found that it did indeed dimerize more quickly than WT (**Fig. 4D**). This finding is consistent with our hypothesis and suggests that the frequency of interactions between the propeptide and the key residues likely plays an essential role in regulating the dimerization rate.

Further statistical analysis confirmed that the dynamics of interactions between the propeptide and the key residues are statistically distinct across the WT, SLOW, and FAST mutant simulations (based on an autocorrelation-robust analysis of three independent replicates for each variant, see **Fig. 5** and the supporting information). Finally, given the capacity for allosteric communication among subunits in a complex like a proteasome (17, 41, 42), we performed a more detailed analysis of the hydrogen bonding dynamics of the individual subunits (**Fig. 6** and **Fig. S8**). This analysis revealed no evidence of any coupling between the hydrogen bonding states of neighboring subunits, suggesting that individual subunits transition between the D+ and D− states independently. Interestingly, we also saw that, in the WT simulations, individual subunits tended to remain in the D− state for around 500-1000μs. The relative stability of this state, along with the fact that multiple subunits might reside in the D− state at any given time, means, on average, that the WT HP spends about 75% of its time in a non-dimerizable state (**Fig. 6A**). In contrast, when the propeptide interacts with the key residues in the SLOW mutant, it forms more hydrogen bonds, making that interaction much more stable (generally longer than the 2.5μs simulations performed here, **Fig. 6B**). Also, more subunits in the SLOW mutant tend to be in the D− state than the WT. As a result, the SLOW mutant essentially never visits a dimerizable state, and, as such, dimerization is essentially entirely prevented. The FAST mutant, however, forms fewer interactions, and as a result, the transitions out of the D− state are much faster than WT (**Fig. 6C**). As a result, the FAST mutant more frequently visits states where all the key residues are available for dimerization, and this likely allows dimerization kinetics to proceed far more rapidly (**Fig. 4D**).

Our results thus strongly suggest that interactions between the propeptide and the key dimerization residues play a critical role in regulating the dimerization rate of the HP in *Re*. Many questions regarding this mechanism still remain unanswered. Perhaps most important is the question of *why* this system has evolved to have a separation of time scales in the first place. The dimerization of two HPs is the rate-limiting step in CP formation, and one would expect there would be an evolutionary pressure to make this step faster. Doing so would likely result in faster response times if the bacterium needed to express more CPs quickly and would almost certainly lead to higher steady-state yields in the bacterial cell (43). Given that even a few mutations of charged residues to alanine in the propeptide can result in faster dimerization, it is *a priori* unclear why evolution has favored a slower-dimerizing variant. One hypothesis for why this separation of time scales exists could be that it acts as a sort of “checkpoint” in the assembly process. Interestingly, in every organism studied to date, there has never been any evidence of dimerization of near-HP intermediates, like, say, α_6_β_6_ or α_6_β_7_, with either similar intermediates or the α_7_β_7_ HP. If an α_6_β_6_ did dimerize with an HP, that could lead to activation of the β subunits and allow untagged proteins access to the proteolytic core, which would lead to uncontrolled proteolysis and could be catastrophic for the cell. Hence, we speculate that a slow dimerization step may have evolved in order to prevent premature activation of incompletely assembled structures.

Another interesting question that arises is how widespread the mechanism is that we have described here. While the proteasome is conserved across all kingdoms of life, the subunit sequences have diverged significantly. In particular, the β propeptide is present on all β subunits but shows comparatively little sequence conservation (21, 44). Even within bacteria, the specific region of the propeptide we have focused on here, Region III, shows very little sequence conservation beyond a clear enrichment in glycine residues (22). Interestingly, many bacterial sequences, including the propeptide from *M. tuberculosis*, have charged residues in Region III that could form contacts with the key residues, but the presence of charged residues in Region III is not universal (**Fig. 1B**). It is thus currently unclear whether this mechanism is conserved within all bacterial proteasomes. Beyond bacteria, the propeptide of the β subunits in archaea is generally very short and is dispensable for assembly (44, 45). Interestingly, however, eukaryotic β subunits often carry long propeptides, even in subunits that have lost their active catalytic site. Thus, while the assembly of the eukaryotic CP is far more complicated than the bacterial CP, involving chaperones like Ump1 that also regulate the dimerization process, it is possible that the mechanism described here may also be deployed in eukaryotic cells as part of an HP assembly checkpoint (8, 21, 22, 46). Further experimental and computational studies are needed to firmly establish and understand the roles played by the propeptide in regulating CP assembly. To date, nearly all studies of proteasome assembly have been experimental. This work demonstrates how biophysical modeling tools like Molecular Dynamics simulation can complement experimental approaches and bring new insights into the molecular mechanisms underlying key steps in the assembly process. This combination of computational modeling and experimental validation will be critical in developing a complete understanding of the biogenesis of the proteasome and other critical molecular machines within the cell.

## Acknowledgments

We would like to thank Prof. Jeroen Roelofs for many helpful discussions on proteasome assembly and for his generous help with generating the FAST mutant described here. We thank Profs. Wonpil Im, Yinglong Miao, Joanna Slusky, Roberto DeGuzman, and Mark Richter for their helpful critique and suggestions regarding this work. The original plasmids containing the PrcA1 and PrcB1 genes from *Re* were a generous gift from W. Baumeister. We would also like to thank the Pittsburgh Supercomputing Center, and specifically Marcela Madrid and Philip Blood, for their help with Anton resource. This work was supported by an NSF grant MCB-1412262 to Eric J. Deeds and Anton allocation grant MCB180087P.

## References

1. Kwon YD, Nagy I, Adams PD, Baumeister W, Jap BK. Crystal structures of the Rhodococcus proteasome with and without its pro-peptides: implications for the role of the pro-peptide in proteasome assembly. J Mol Biol. 2004;335(1):233–45.

2. Sharon M, Witt S, Glasmacher E, Baumeister W, Robinson CV. Mass spectrometry reveals the missing links in the assembly pathway of the bacterial 20 S proteasome. J Biol Chem. 2007;282(25):18448–57.

3. Baumeister W, Walz J, Zuhl F, Seemuller E. The proteasome: paradigm of a self-compartmentalizing protease. Cell. 1998;92(3):367–80.

4. Witt S, Kwon YD, Sharon M, Felderer K, Beuttler M, Robinson CV, et al. Proteasome assembly triggers a switch required for active-site maturation. Structure. 2006;14(7):1179–88.

5. Zwickl P. The 20S proteasome. Curr Top Microbiol Immunol. 2002;268:23–41.

6. Jastrab JB, Darwin KH. Bacterial Proteasomes. Annu Rev Microbiol. 2015;69:109–27.

7. Zuhl F, Seemuller E, Golbik R, Baumeister W. Dissecting the assembly pathway of the 20S proteasome. FEBS Lett. 1997;418(1-2):189–94.

8. Shigeo Murata HYaKT. Molecular mechanisms of proteasome assembly. Nature reviews. 2009;10.

9. Cheng Y, Pieters J. Novel proteasome inhibitors as potential drugs to combat tuberculosis. J Mol Cell Biol. 2010;2(4):173–5.

10. Lin G, Li D, Chidawanyika T, Nathan C, Li H. Fellutamide B is a potent inhibitor of the Mycobacterium tuberculosis proteasome. Arch Biochem Biophys. 2010;501(2):214–20.

11. Field-Smith A, Morgan GJ, Davies FE. Bortezomib (Velcadetrade mark) in the Treatment of Multiple Myeloma. Ther Clin Risk Manag. 2006;2(3):271–9.

12. Kane RC, Bross PF, Farrell AT, Pazdur R. Velcade: U.S. FDA approval for the treatment of multiple myeloma progressing on prior therapy. Oncologist. 2003;8(6):508–13.

13. Stock D, Ditzel L, Baumeister W, Huber R, Lowe J. Catalytic mechanism of the 20S proteasome of Thermoplasma acidophilum revealed by X-ray crystallography. Cold Spring Harb Symp Quant Biol. 1995;60:525–32.

14. Panfair D, Ramamurthy A, Kusmierczyk AR. Alpha-ring Independent Assembly of the 20S Proteasome. Sci Rep. 2015;5:13130.

15. Tamura T, Nagy I, Lupas A, Lottspeich F, Cejka Z, Schoofs G, et al. The first characterization of a eubacterial proteasome: the 20S complex of Rhodococcus. Curr Biol. 1995;5(7):766–74.

16. Wilson HL, Ou MS, Aldrich HC, Maupin-Furlow J. Biochemical and physical properties of the Methanococcus jannaschii 20S proteasome and PAN, a homolog of the ATPase (Rpt) subunits of the eucaryal 26S proteasome. J Bacteriol. 2000;182(6):1680–92.

17. Wani PS, Rowland MA, Ondracek A, Deeds EJ, Roelofs J. Maturation of the proteasome core particle induces an affinity switch that controls regulatory particle association. Nat Commun. 2015;6:6384.

18. Seemuller E, Lupas A, Baumeister W. Autocatalytic processing of the 20S proteasome. Nature. 1996;382(6590):468–71.

19. Maupin-Furlow JA, Aldrich HC, Ferry JG. Biochemical characterization of the 20S proteasome from the methanoarchaeon Methanosarcina thermophila. J Bacteriol. 1998;180(6):1480–7.

20. Brewer JW, Corley RB. Late events in assembly determine the polymeric structure and biological activity of secretory IgM. Mol Immunol. 1997;34(4):323–31.

21. Seemller E, Zwickl P, Baumeister W. Self-processing of Subunits of the Proteasome. The Enzymes. 2001;22:335–71.

22. Suppahia A, Itagi P, Burris A, Kim FMG, Vontz A, Kante A, et al. Cooperativity in Proteasome Core Particle Maturation. iScience. 2020;23(5):101090.

23. Zwickl P, Kleinz J, Baumeister W. Critical elements in proteasome assembly. Nat Struct Biol. 1994;1(11):765–70.

24. Mayr J, Seemuller E, Muller SA, Engel A, Baumeister W. Late events in the assembly of 20S proteasomes. J Struct Biol. 1998;124(2-3):179–88.

25. Shaw DE, Grossman JP, Bank JA, Batson B, Butts JA, Chao JC, et al., editors. Anton 2: Raising the Bar for Performance and Programmability in a Special-Purpose Molecular Dynamics Supercomputer. SC ’14: Proceedings of the International Conference for High Performance Computing, Networking, Storage and Analysis; 2014 16-21 Nov. 2014.

26. Schrodinger. The PyMOL Molecular Graphics System, Version 2.0 Schrödinger, LLC. 2015.

27. Raman S, Vernon R, Thompson J, Tyka M, Sadreyev R, Pei J, et al. Structure prediction for CASP8 with all-atom refinement using Rosetta. Proteins. 2009;77 Suppl 9(0 9):89–99.

28. Song Y, DiMaio F, Wang RY, Kim D, Miles C, Brunette T, et al. High-resolution comparative modeling with RosettaCM. Structure. 2013;21(10):1735–42.

29. Jo S, Kim T, Iyer VG, Im W. CHARMM-GUI: a web-based graphical user interface for CHARMM. J Comput Chem. 2008;29(11):1859–65.

30. Huang J, Rauscher S, Nawrocki G, Ran T, Feig M, de Groot BL, et al. CHARMM36m: an improved force field for folded and intrinsically disordered proteins. Nature Methods. 2017;14(1):71–3.

31. Lee J, Cheng X, Swails JM, Yeom MS, Eastman PK, Lemkul JA, et al. CHARMM-GUI Input Generator for NAMD, GROMACS, AMBER, OpenMM, and CHARMM/OpenMM Simulations Using the CHARMM36 Additive Force Field. J Chem Theory Comput. 2016;12(1):405–13.

32. Case DA, Cheatham TE, 3rd, Darden T, Gohlke H, Luo R, Merz KM, Jr., et al. The Amber biomolecular simulation programs. J Comput Chem. 2005;26(16):1668–88.

33. Salomon-Ferrer R, Götz AW, Poole D, Le Grand S, Walker RC. Routine Microsecond Molecular Dynamics Simulations with AMBER on GPUs. 2. Explicit Solvent Particle Mesh Ewald. Journal of Chemical Theory and Computation. 2013;9(9):3878–88.

34. Lippert RA, Predescu C, Ierardi DJ, Mackenzie KM, Eastwood MP, Dror RO, et al. Accurate and efficient integration for molecular dynamics simulations at constant temperature and pressure. J Chem Phys. 2013;139(16):164106.

35. Newey W, West K. Automatic Lag Selection in Covariance Matrix Estimation. Review of Economic Studies. 1994;61(4):631–53.

36. Newey W, West K. A Simple, Positive Semi-definite, Heteroskedasticity and Autocorrelation Consistent Covariance Matrix. Econometrica. 1987;55(3):703–08.

37. Andrews D. Heteroskedasticity and Autocorrelation Consistent Covariance Matrix Estimation. Econometrica. 1991;59(3):817–58.

38. Zeileis A. Econometric Computing with HC and HAC Covariance Matrix Estimators. Journal of Statistical Software. 2004;11(10):1–17.

39. RDC. T. R: A language and environment for statistical computing. Vienna, Austria 2010.

40. Cleveland WS. Robust Locally Weighted Regression and Smoothing Scatterplots. Journal of the American Statistical Association. 1979;74(368):829–36.

41. Bray D, Duke T. Conformational spread: the propagation of allosteric states in large multiprotein complexes. Annu Rev Biophys Biomol Struct. 2004;33:53–73.

42. Bray D, Levin MD, Lipkow K. The chemotactic behavior of computer-based surrogate bacteria. Curr Biol. 2007;17(1):12–9.

43. Deeds EJ, Bachman JA, Fontana W. Optimizing ring assembly reveals the strength of weak interactions. Proc Natl Acad Sci U S A. 2012;109(7):2348–53.

44. Marques AJ, Palanimurugan R, Matias AC, Ramos PC, Dohmen RJ. Catalytic mechanism and assembly of the proteasome. Chem Rev. 2009;109(4):1509–36.

45. Zwickl P, Grziwa A, Puhler G, Dahlmann B, Lottspeich F, Baumeister W. Primary structure of the Thermoplasma proteasome and its implications for the structure, function, and evolution of the multicatalytic proteinase. Biochemistry. 1992;31(4):964–72.

46. Zühl F, Tamura T, Dolenc I, Cejka Z, Nagy I, De Mot R, et al. Subunit topology of the Rhodococcus proteasome. FEBS Lett. 1997;400(1):83–90.

