## Supplemental Information for "Understanding the Separation of Timescales in *Rhodococcus erythropolis* Proteasome Core Particle Assembly"

Biophysical Journal

#### S.1 Propeptide comes out of the Half Proteasome( HP) barrel in WT simulations

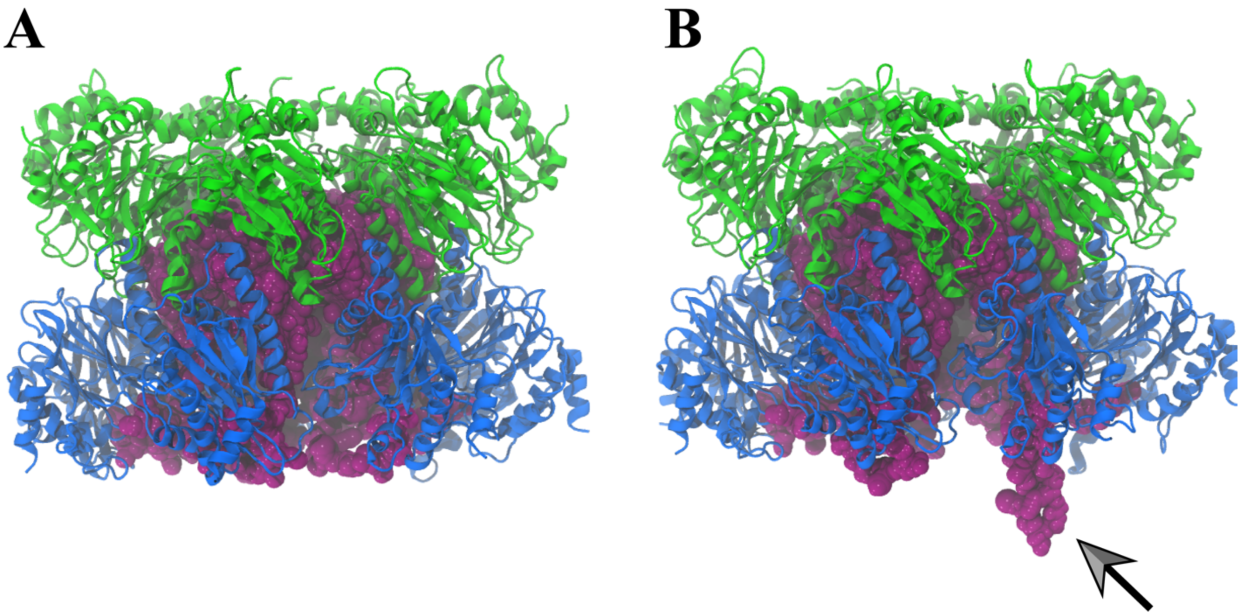


**Figure S.1**. Cartoon representation of the WT HP from Molecular Dynamics simulations. The α subunits are shown in green, β subunits in blue and the β propeptides in the purple spheres. (A) The WT HP at 996 ns. (B) WT HP at 1240.8 ns. The arrow serves to highlight the protruding propeptide. Both images were rendered using VMD. (1)

### S.2 Root Mean Square Fluctuation (Å) for the propeptide in WT, SLOW and FAST simulations

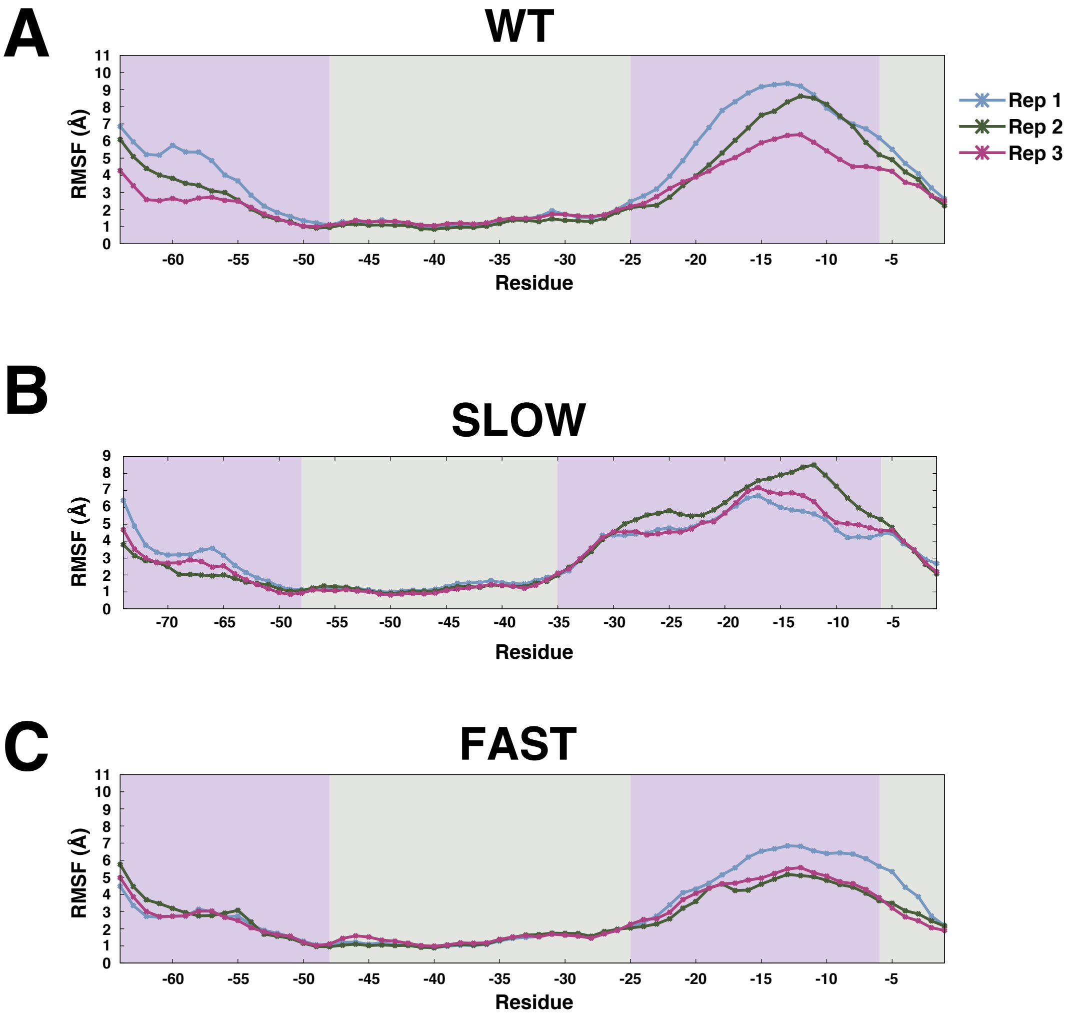


**Figure S.2.** Root Mean-Square Fluctuations (RMSF) of the β propeptide in WT, SLOW and FAST HPs. The panels show the average RMSF after 2.5 μs for the backbone atoms of each residue of the β propeptides in (A) WT, (B) SLOW and (C) FAST HP for three independent Molecular Dynamics simulations. For all panels, electron density of the residues in crystal structure is depicted by the colored bars; missing electron density residues (purple) and residues with electron density (grey). The missing electron density residues were modeled by Rosetta. The RMSF values measured for Replicate Rep. 1 is shown by the blue curve, Rep. 2 by the green and Rep. 3 by the pink colors.

### S.3 Potential energy profiles of the Anton simulations

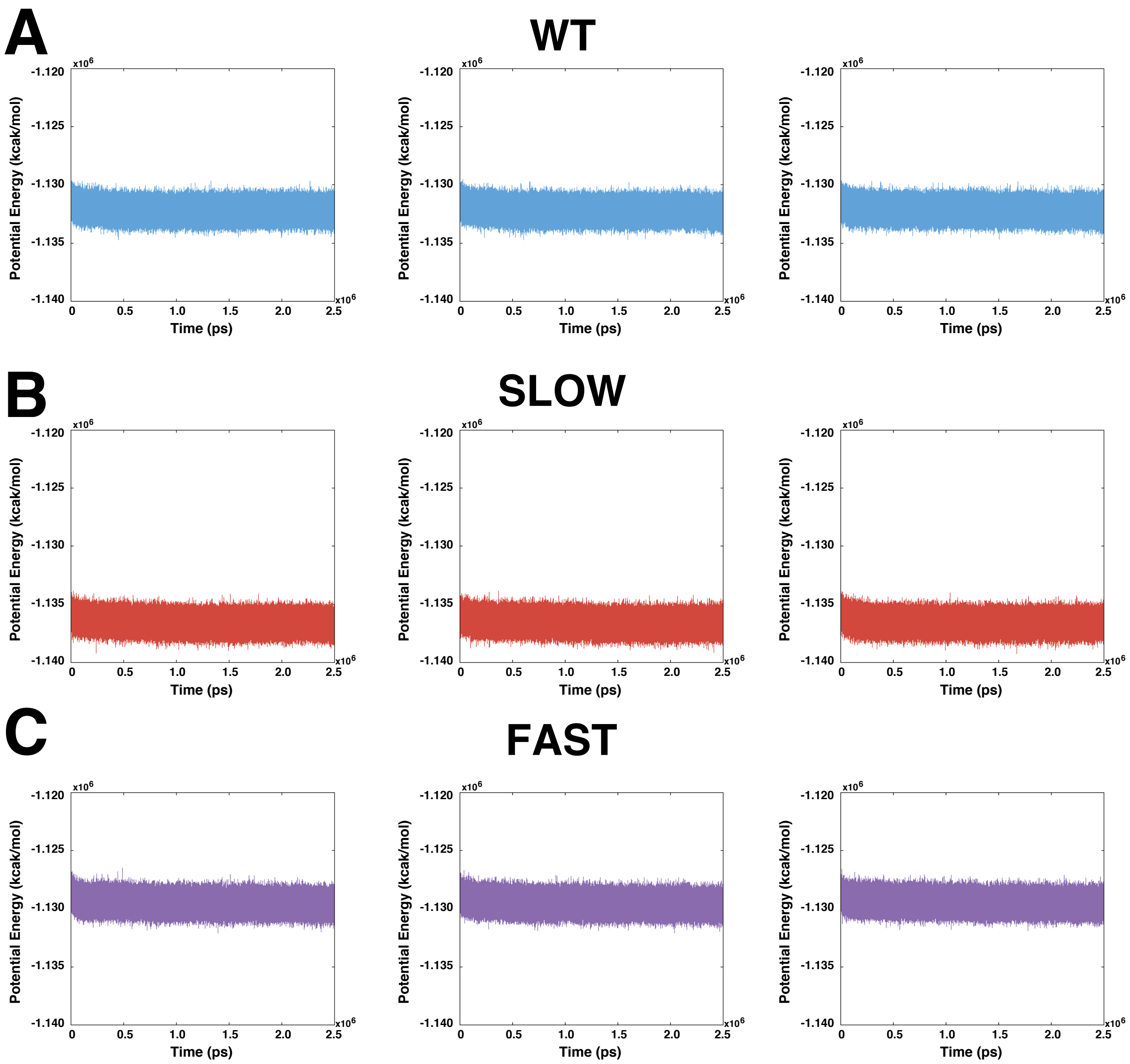


**Figure S.3**. Potential energy as a function of timer for each simulation. The potential energy of each 2.5 μs simulation on Anton with (A) WT, (B) SLOW and (C) FAST HPs. Plots show the potential energy for Rep. 1 (left), Rep. 2 (middle) and Rep. 3 (right) separately for each HP type. Every simulation took about 500 ns to converge.

### S.4 RMSD (without propeptide) of all backbone atoms as a function of time

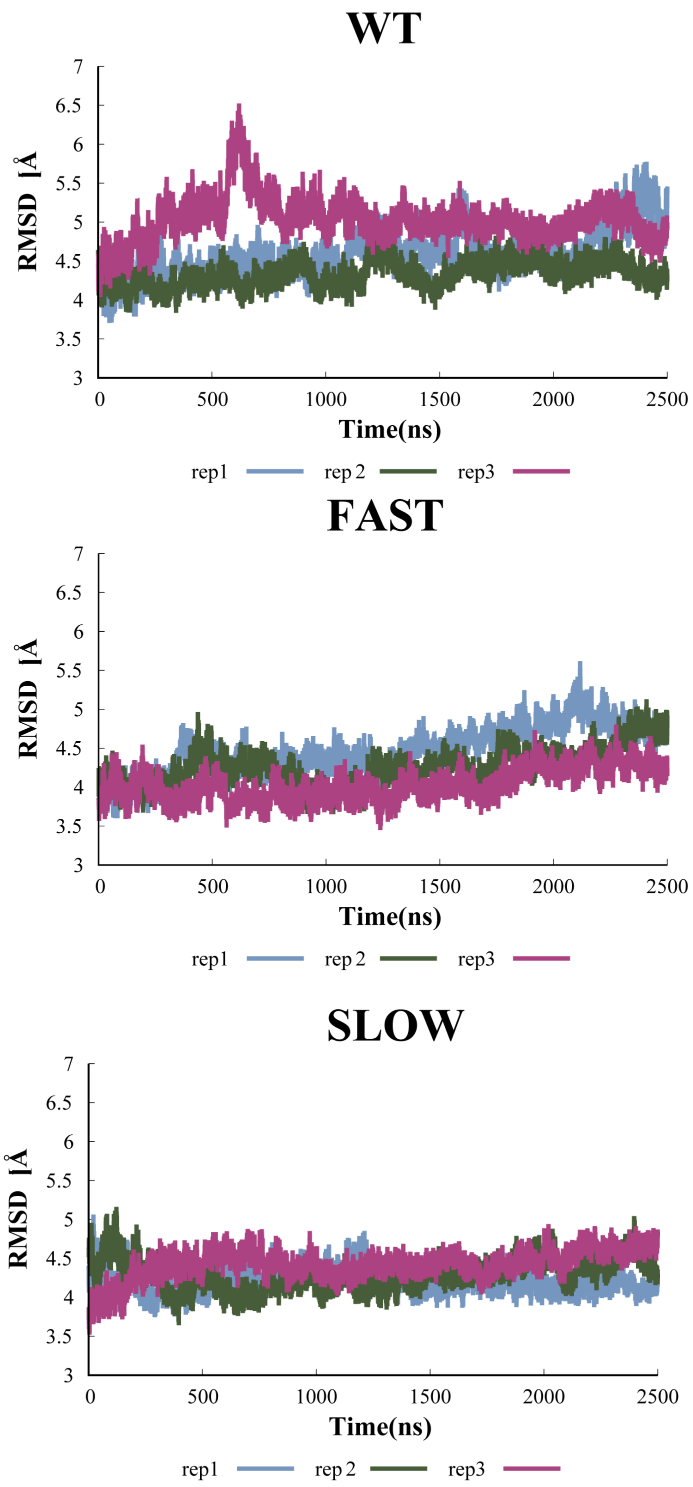


**Figure S.4**. RMSD for HP without the propeptide. Panels show the average RMSD for all seven β subunits of the HP without including the propeptide residues in the RMSD calculations at each time point for (A) WT, (B) SLOW and (C) FAST HPs. For comparison, all three replicates (blue, green and pink) are shown on the same axes.

**S.5 Locally Weighted Scatterplot Smoothing (LOWESS) plots of hydrogen bonds between propeptide and key residues**

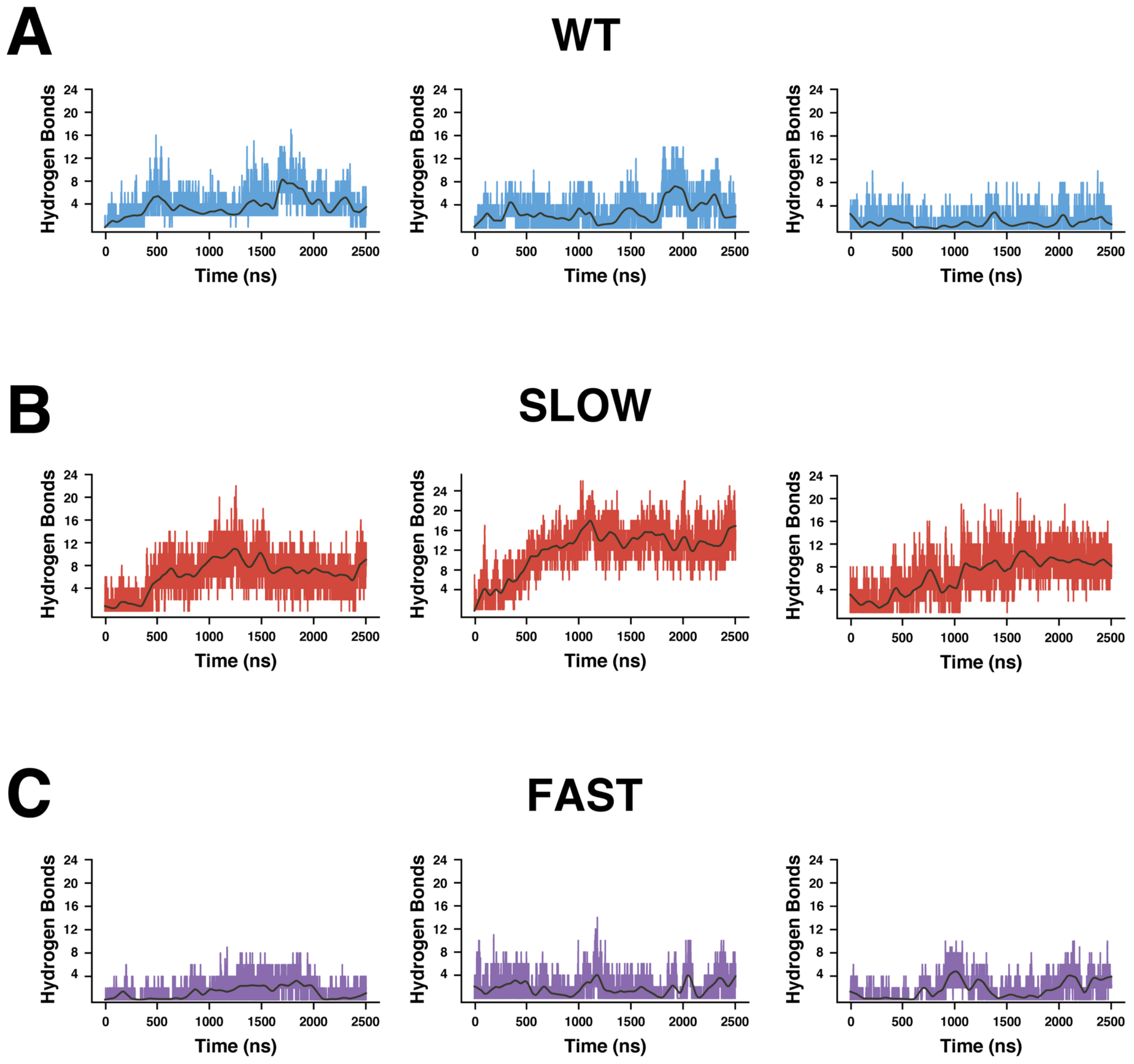


**Figure S.5**. LOWESS plots for HPs. Plots show the number of hydrogen bonds formed by the propeptide of (A) WT, (B) SLOW and (C) FAST HPs at each time point. Plots show the hydrogen bonds for Rep. 1 (left), Rep. 2 (middle) and Rep. 3 (right) separately for each HP type. Gray line represents the non-parametric LOWESS fit.

### S.6 Hydrogen bond distributions for all replicates of WT, SLOW and FAST.

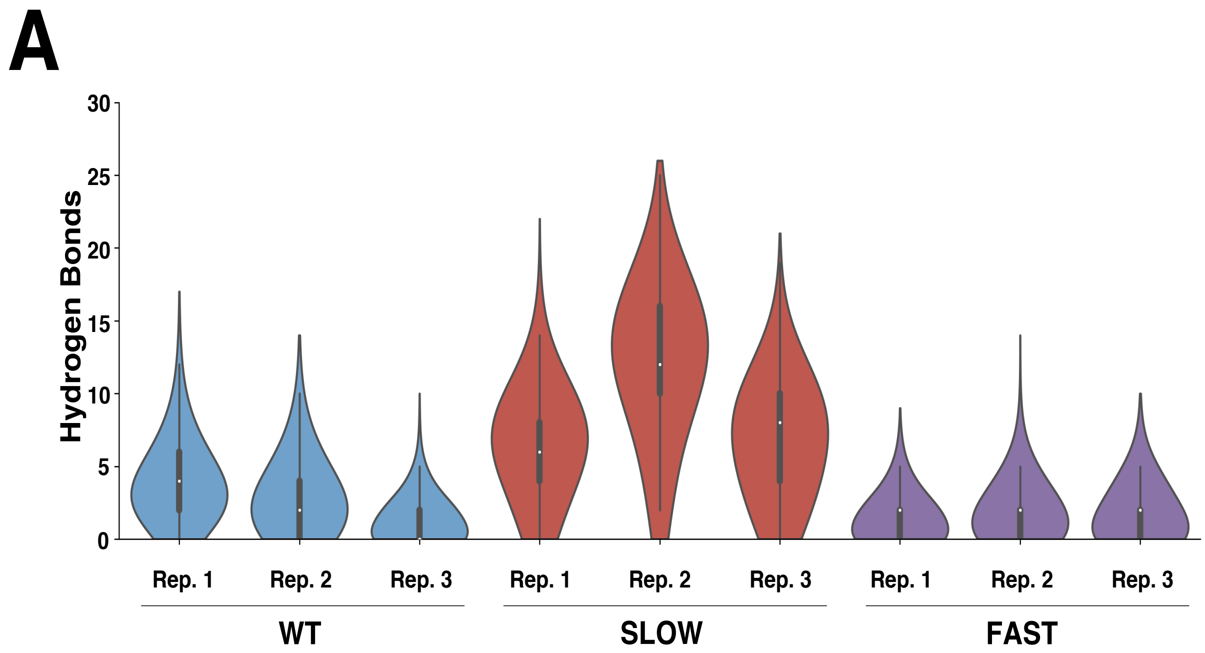


**Figure S.6.** Violin plots of hydrogen bonds formed by propeptide. Violin plots showing the total number of hydrogen bonds formed between the propeptide and key residues at the HP dimerization interface for each replicate of WT (blue), SLOW (red) and FAST (purple) HP.

### S.7 Total number of hydrogen bonds formed between the propeptide residues -9 and -12 and the key residues for WT and FAST Half-Proteasomes (HPs).

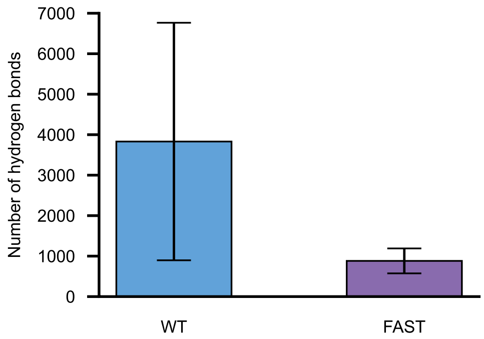


**Figure S.7:** The WT and FAST total hydrogen bonds made specifically by propeptide residues -9(GLU) and -12(ASP), which are mutated to Alanine in FAST mutant. The FAST HPs formed fewer hydrogen bonds compared to the WT HPs.

### S.8 Kymographs for all replicates of WT, SLOW and FAST

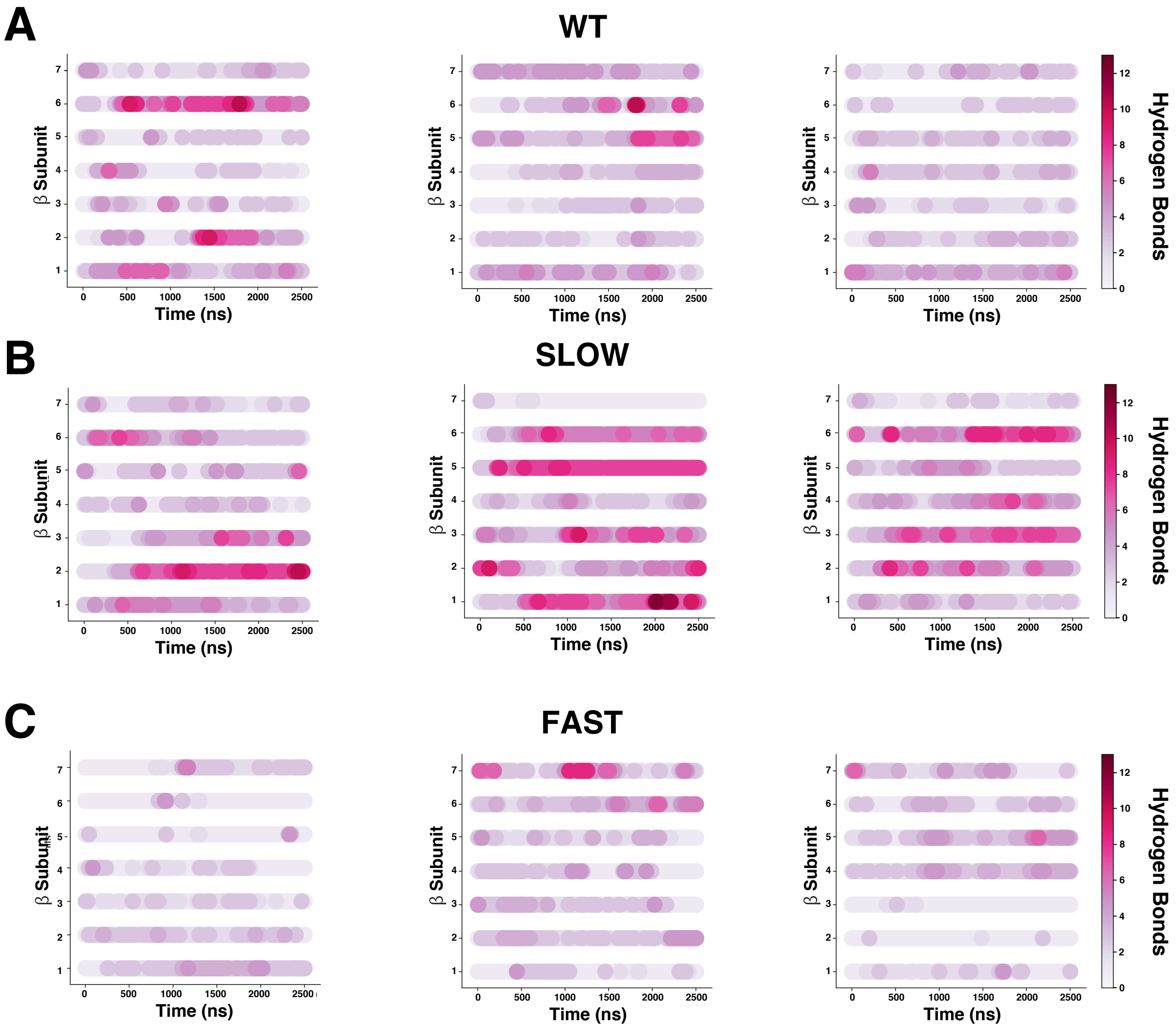


**Figure S.8**. Kymographs for each HP type. The kymographs show the total number of hydrogen bonds formed between each β subunits and the key resides at each time point for (A) WT, (B) SLOW and (C) FAST HPs. Plots show the hydrogen bonds for Rep. 1 (left), Rep. 2 (middle) and Rep. 3 (right) separately for each HP type.

### S.9 Complete results for the categorical regression analysis

#### S.9.1. Model 1 statistical *p-values:* one intercept and one slope

| Simulation | Intercept<br>Intercept $(\boldsymbol{\beta_0})$ *p-value* | Slope<br>Slope $(\boldsymbol{\beta_1})$ *p-value* |
|---|---|---|
| WT-rep1 | $< 2.2\times10^{-16}$ | $1.65\times10^{-5}$ |
| WT-rep2 | $2.0\times10^{-4}$ | $< 2.2\times10^{-16}$ |
| WT-rep3 | $2.7\times10^{-4}$ | $1.73\times10^{-12}$ |
| SLOW-rep1 | $< 2.2\times10^{-16}$ | $7.6\times10^{-4}$ |
| SLOW-rep2 | $< 2.2\times10^{-16}$ | $2.05\times10^{-5}$ |
| SLOW-rep3 | $< 2.2e\times10^{-16}$ | $< 2.2\times10^{-16}$ |
| FAST-rep1 | $< 2\times10^{-16}$ | 0.2072 |
| FAST-rep2 | $6.77\times10^{-9}$ | $3.1\times10^{-4}$ |
| FAST-rep3 | $1.37\times10^{-6}$ | $6.94\times10^{-9}$ |

**Table S.9.1:** Table for categorical regression p-values for the intercept and slope using Newey-West estimator fits for number of hydrogen bonds as a function of time (500ns to 2.5 μs). The simulations whose slope *p-values* is not significant is highlighted in grey.

The first model we considered is of the form $y = \beta_0 + \beta_1 t + \varepsilon$ where $y$ is the number of hydrogen bonds between β propeptide residues and key residues at the HP dimerization interface, $t$ is the simulation time, $\beta_0$ is the intercept, $\beta_1$ is the slope, and $\varepsilon$ is the error term. This basic model essentially asks if there is a significant trend in the number of hydrogen bonds formed between the propeptide residues and the key residues over time. To account for both the possibility of heteroscedasticity and the inherent autocorrelations in individual MD simulations, we used the robust Newey-West estimator to compute p-values (2, 3). As shown in Table **S.9.1**, the p-values are, by and large, highly significant for this regression. This suggests that most replicates, except for the first replicate of the FAST mutant, show a statistically significant increasing trend in the total number of hydrogen bonds over time. While these results are suggestive that total numbers of hydrogen bonds might increase if we were able to run simulations for longer timescales, the primary purpose of these regressions is to serve as a basis for the further statistical analyses presented below.

#### S.9.2 Model 2 statistical *p-values:* two intercepts and one slope

| Wild-Type (WT) | Mutant | WT-Intercept ($\beta_0$) *p-value* | WT & Mutant-Slope ($\beta_1$) *p-value* | Mutant-Intercept ($\beta_2$) *p-value* |
|---|---|---|---|---|
| WT-rep1 | SLOW-rep1 | $3.92\times10^{-16}$ | 0.2527 | $< 2.2\times10^{-16}$ |
| | SLOWrep2 | $< 2.2\times10^{-16}$ | $1.98\times10^{-5}$ | $< 2.2\times10^{-16}$ |
| | SLOW-rep3 | $3.92\times10^{-16}$ | $< 2.2\times10^{-16}$ | $< 2.2\times10^{-16}$ |
| | FAST-rep1 | $< 2.2\times10^{-16}$ | $3.86\times10^{-5}$ | $9.70\times10^{-8}$ |
| | FAST-rep2 | $< 2.2\times10^{-16}$ | $9.70\times10^{-8}$ | $< 2.2\times10^{-16}$ |
| | FAST-rep3 | $< 2.2\times10^{-16}$ | $5.66\times10^{-11}$ | $< 2.2\times10^{-16}$ |
| WT-rep2 | SLOW-rep1 | $< 2.2\times10^{-16}$ | $1.23\times10^{-6}$ | $< 2.2\times10^{-16}$ |
| | SLOW-rep2 | $1.15\times10^{-3}$ | $< 2.2\times10^{-16}$ | $< 2.2\times10^{-16}$ |
| | SLOW-rep3 | 0.3865 | $< 2.2\times10^{-16}$ | $< 2.2\times10^{-16}$ |
| | FAST-rep1 | $< 2.2\times10^{-16}$ | $< 2.2\times10^{-16}$ | $< 2.2\times10^{-16}$ |
| | FAST-rep2 | $< 2.2\times10^{-16}$ | $< 2.2\times10^{-16}$ | $< 2.2\times10^{-16}$ |
| | FAST-rep3 | $< 2.2\times10^{-16}$ | $< 2.2\times10^{-16}$ | $6.14\times10^{-13}$ |
| WT-rep3 | SLOW-rep1 | $2.87\times10^{-14}$ | 0.3975 | $< 2.2\times10^{-16}$ |
| | SLOW-rep2 | 0.3425 | $6.04\times10^{-10}$ | $< 2.2\times10^{-16}$ |
| | SLOW-rep3 | $7.44\times10^{-14}$ | $< 2.2\times10^{-16}$ | $< 2.2\times10^{-16}$ |
| | FAST-rep1 | $1.67\times10^{-10}$ | $3.79\times10^{-7}$ | $7.75\times10^{-7}$ |
| | FAST-rep2 | $3.48\times10^{-4}$ | $8.59\times10^{-11}$ | $2.59\times10^{-7}$ |
| | FAST-rep3 | 0.0479 | $3.614\times10^{-16}$ | $< 2.2\times10^{-16}$ |

**Table S.9.2**: Table for categorical regression *p-values* for the intercepts and slope from Newey-West estimator fits for the number of hydrogen bonds as a function of time (500ns to 2.5 μs). The simulations whose slope *p-values* are not significant are highlighted in grey, and the *p-values* where the intercept is not significant are in red.

The second model we considered is aimed at *comparing* the WT simulations to the mutant simulations. As mentioned in the main text, performing a standard statistical test on the hydrogen bond distributions (Fig. S.6) will not work in this case, because the samples that make up this distribution are highly autocorrelated and thus violate the standard "Independent and Identically Distributed" assumption made by essentially every statistical test for a difference in means. To overcome this problem, we employed a categorical regression using the robust Newey-West estimator to account for autocorrelations. The model in this case is $y = \beta_0 + \beta_1 t + \beta_2 C + \varepsilon$, where $y$ is the number of hydrogen bonds between propeptide residues and the key residues at HP dimerization interface, $t$ is the simulation time, $C$ is a categorical variable and the $\beta$'s are the regression coefficients. The idea here is that we combine the results from one WT simulation and one mutant simulation into the same regression. At any given time in this case, there are actually two possible values of $y$; one from the WT simulation and one from the mutant. The categorical variable $C$ distinguishes between the WT and mutant simulations: $C = 0$ for WT data points and $C = 1$ for the mutant. Note that, in this model, if the coefficient $\beta_2 = 0$, then $y = \beta_0 + \beta_1 t + \varepsilon$; in other words, the slope and the intercept would be *the same* for the WT and mutant cases. When $\beta_2 \neq 0$, then we have essentially two different regressions happening at the same time: $y = \beta_0 + \beta_1 t + \varepsilon$ for WT data points with $C = 0$ and $y = \beta_0 + \beta_1 t + \beta_2 + \varepsilon$ for mutant data points with $C = 1$. This essentially means that, when $\beta_2 \neq 0$, the WT and mutant regressions have different intercepts but the same slope.

Recall that the p-value for any given coefficient in a regression like this represents the probability that we would observe a coefficient value at least as large under the null hypothesis that the coefficient in question was actually 0. The p-value represents our statistical confidence in a non-zero regression coefficient. So, a significant p-value for $\beta_2$ indicates that the intercepts for the WT and mutant cases are statistically different. The results for this regression are shown in Table **S.9.2**. Interestingly, the only non-significant results are for the coefficients $\beta_0$ and $\beta_1$; the results for $\beta_2$ are all highly significant. This indicates that the intercept value for the mutant is always distinct from the WT in this model. Note that, in this case, we have forced the mutant and WT cases to have the same slope.

**S9.3 Model 3 *p-values:* two intercepts and two slopes=**

| Wild-Type (WT) | Mutant | WT- Intercept ($\beta_0$) *p-value* | WT-Slope ($\beta_1$) *p-value* | Mutant-Intercept ($\beta_2$) *p-value* | Mutant-Slope ($\beta_3$) *p-value* |
|---|---|---|---|---|---|
| WT rep 1 | SLOW-rep1 | $< 2.2\times10^{-16}$ | $5.49\times10^{-5}$ | $< 2.2\times10^{-16}$ | $3.65\times10^{-6}$ |
| | SLOW-rep2 | $< 2.2\times10^{-16}$ | $5.05\times10^{-5}$ | $< 2.2\times10^{-16}$ | 0.5556 |
| | SLOW-rep3 | $< 2.2\times10^{-16}$ | $5.47\times10^{-5}$ | 0.1767 | $< 2.2\times10^{-16}$ |
| | FAST-rep1 | $< 2.2\times10^{-16}$ | $5.50\times10^{-5}$ | $1.142\times10^{-10}$ | 0.0053 |
| | FAST-rep2 | $< 2.2\times10^{-16}$ | $6.50\times10^{-5}$ | $1.410\times10^{-11}$ | 0.3192 |
| | FAST-rep3 | $< 2.2\times10^{-16}$ | $< 5.09\times10^{-5}$ | $4.706\times10^{-12}$ | 0.8151 |
| WT-rep2 | SLOW-rep1 | 0.0004 | $< 2.2\times10^{-16}$ | $< 2.2\times10^{-16}$ | $< 2.2\times10^{-16}$ |
| | SLOW-rep2 | 0.0003 | $< 2.2\times10^{-16}$ | $< 2.2\times10^{-16}$ | 0.0072 |
| | SLOW-rep3 | 0.0004 | $< 2.2\times10^{-16}$ | $4.71\times10^{-16}$ | $6.28\times10^{-7}$ |
| | FAST-rep1 | 0.0004 | $< 2.2\times10^{-16}$ | 0.0104 | $9.26\times10^{-13}$ |
| | FAST-rep2 | 0.0004 | $< 2.2\times10^{-16}$ | 0.2020 | $5.27\times10^{-7}$ |
| | FAST-rep3 | 0.0003 | $< 2.2\times10^{-16}$ | 0.3905 | 0.0001 |
| WT-rep3 | SLOW-rep1 | 0.0043 | $1.86\times10^{-11}$ | $< 2.2\times10^{-16}$ | $1.12\times10^{-7}$ |
| | SLOW-rep2 | 0.0041 | $1.57\times10^{-11}$ | $< 2.2\times10^{-16}$ | 0.2489 |
| | SLOW-rep3 | 0.0043 | $1.85\times10^{-11}$ | $< 2.2\times10^{-16}$ | $< 2.2\times10^{-16}$ |
| | FAST-rep1 | 0.0043 | $1.90\times10^{-11}$ | $9.07\times10^{-8}$ | 0.0009 |
| | FAST-rep2 | 0.0046 | $2.68\times10^{-11}$ | 0.0013 | 0.4784 |
| | FAST-rep3 | 0.0042 | $1.63\times10^{-11}$ | 0.0099 | 0.3358 |

**Figure S.9.3**: Table for categorical regression *p-values* for the intercepts and slopes estimated from Newey-West estimators fits for the number of hydrogen bonds as a function of time (500ns to 2.5 μs). The simulations where slope *p-values* are not significant are highlighted in grey background, , and the *p-values* where intercept are not significant is in red.

The final model we consider in this analysis allows the WT and mutant to have both a different slope and a different intercept. The model takes the form $y = \beta_0 + \beta_1 t + \beta_2 C + \beta_3 Ct + \varepsilon$ where $y$ is the number of hydrogen bonds between propeptide residues and the set of key residues at HP dimerization interface, $t$ is the simulation time, $C$ is the categorical variable as described in the previous section, $\varepsilon$ is the error term and the $\beta$'s are the regression coefficients. Similar to the discussion above, significant p-values for either $\beta_2$ or $\beta_3$ indicate statistical support for a different intercept or slope between the WT and mutant replicate being compared. Note that, in this model, for every comparison between the WT and mutant replicates, either the intercept or slope term is statistically significant. This strongly suggests that the behavior of the WT simulations is different from those of the various mutants, supporting the idea that the formation of hydrogen bonds between the propeptide and the key residues follows significantly different trends in the SLOW and FAST simulations compared to the WT.

### S. 10 MD simulations systems details

| Simulation | Box size (Å) | Number of atoms | Number of water molecules | Ions |
|---|---|---|---|---|
| WT | 155 ×155 ×155 | 352979 | 298725 | $Na^+$ = 275<br>$Cl^-$ = 184 |
| SLOW | 155 ×155 ×155 | 353567 | 298515 | $Na^+$ = 275<br>$Cl^-$ = 184 |
| FAST | 155 ×155×155 | 345648 | 291465 | $Na^+$ = 257<br>$Cl^-$ = 180 |

**Table A. 10:** System properties and details of the HP, FAST, and SLOW simulations. All the simulations are run in a rectangular water box with 15 Å water on each side of the protein.

### S. 11. Additional MD simulations details.

All simulations were performed using the CHARMM36 force field and explicit water was modeled with the CHARMM version(4, 5) of the TIP3P water model (6). All the simulations are explicit solvent using TIP3P water model with CHARMM36m version(7). In the Anton2 simulations, integration was carried out using the Multigrator algorithm (8) with a 2.5 fs time step. For the RESPA scheme every second time step was used for long range interactions. Pressure was controlled using the Martyna-Tobias-Klein (MTK) barostat (9) with an interval length of 480 ps. The temperature was maintained by the Nose-Hoover thermostat (10) with an interval length of 24 ps. A relaxation time of τ=0.041667 ps was used for both the barostat and thermostat.

### References


1. Humphrey W, Dalke A, Schulten K. VMD: visual molecular dynamics. J Mol Graph. 1996;14(1):33-8, 27-8.
2. Newey W, West K. Automatic Lag Selection in Covariance Matrix Estimation. Review of Economic Studies. 1994;61(4):631-53.
3. Newey W, West K. A Simple, Positive Semi-definite, Heteroskedasticity and Autocorrelation Consistent Covariance Matrix. Econometrica. 1987;55(3):703-08.
4. Best RB, Zhu X, Shim J, Lopes PE, Mittal J, Feig M, et al. Optimization of the additive CHARMM all-atom protein force field targeting improved sampling of the backbone φ, ψ and side-chain χ(1) and χ(2) dihedral angles. J Chem Theory Comput. 2012;8(9):3257-73.
5. MacKerell AD, Bashford D, Bellott M, Dunbrack RL, Evanseck JD, Field MJ, et al. All-atom empirical potential for molecular modeling and dynamics studies of proteins. J Phys Chem B. 1998;102(18):3586-616.
6. Jorgensen WL, Chandrasekhar J, Madura JD, Impey RW, Klein ML. Comparison of simple potential functions for simulating liquid water. Journal of Chemical Physics. 1983;79:926.
7. Huang J, Rauscher S, Nawrocki G, Ran T, Feig M, de Groot BL, et al. CHARMM36m: an improved force field for folded and intrinsically disordered proteins. Nature Methods. 2017;14(1):71-3.
8. Lippert RA, Predescu C, Ierardi DJ, Mackenzie KM, Eastwood MP, Dror RO, et al. Accurate and efficient integration for molecular dynamics simulations at constant temperature and pressure. The Journal of Chemical Physics. 2013;139(16):164106.
9. Martyna GJ, Tobias DJ, Klein ML. Constant pressure molecular dynamics algorithms. The Journal of Chemical Physics. 1994;101(5):4177-89.
10. Nosé S. A molecular dynamics method for simulations in the canonical ensemble. Molecular Physics. 1984;52(2):255-68.
